# Ser/Thr phospho-regulation by PknB and Stp mediates bacterial quiescence and antibiotic persistence in *Staphylococcus aureus*

**DOI:** 10.1101/2021.05.06.442895

**Authors:** Markus Huemer, Srikanth Mairpady Shambat, Sandro Pereira, Lies Van Gestel, Judith Bergada-Pijuan, Alejandro Gómez-Mejia, Chun-Chi Chang, Clement Vulin, Julian Bär, Jonathan Dworkin, Annelies S. Zinkernagel

**Affiliations:** Department of Infectious Diseases, University Hospital Zurich, University of Zurich, Zurich, Switzerland; University of Hasselt (UHasselt), Hasselt, Belgium; Department of Microbiology and Immunology, College of Physicians and Surgeons, Columbia University, New York, USA

**Keywords:** *Staphylococcus aureus*, antibiotic persistence, Stp, PknB, phosphoproteome, quiescence

## Abstract

*Staphylococcus aureus* colonizes 30 to 50% of healthy adults and can cause a variety of diseases, ranging from superficial to life-threatening invasive infections such as bacteraemia and endocarditis. Often, these infections are chronic and difficult-to-treat despite adequate antibiotic therapy. Most antibiotics act on metabolically active bacteria in order to eradicate them. Thus, bacteria with minimized energy consumption resulting in metabolic quiescence, have increased tolerance to antibiotics. The most energy intensive process in cells – protein synthesis – is attenuated in bacteria entering into quiescence. Eukaryote-like serine/threonine kinases (STKs) and phosphatases (STPs) can fine-tune essential cellular processes, thereby enabling bacteria to quickly respond to environmental changes and to modulate quiescence. Here, we show that deletion of the only annotated functional STP, named Stp, in *S. aureus* leads to increased bacterial lag-phase and phenotypic heterogeneity under different stress challenges, including acidic pH, intracellular milieu and *in vivo* abscess environment. This growth delay was associated with reduced intracellular ATP levels and increased antibiotic persistence. Using phosphopeptide enrichment and mass spectrometry-based proteomics, we identified possible targets of Ser/Thr phosphorylation that regulate cellular processes and bacterial growth, such as ribosomal proteins including the essential translation elongation factor EF-G. Finally, we show that acid stress leads to a reduced translational activity in the *stp* deletion mutant indicating metabolic quiescence correlating with increased antibiotic persistence.

**One-sentence summary:** Phospho-regulation mediates quiescence and antibiotic persistence in *Staphylococcus aureus*.

## Main

The Gram-positive bacterium *Staphylococcus aureus* intermittently or permanently colonizes 30-50% of the human population ^1,2^. Although *S. aureus* is part of the normal skin microbiota, it is a leading cause of a variety of nosocomial and community-acquired infections ^1^. Besides skin and soft tissue infections (SSTI), *S. aureus* can cause severe and life-threatening, invasive infections such as endocarditis, osteomyelitis and bacteraemia ^1^. Increasing levels of resistance against multiple antibacterial drugs, including penicillin, methicillin and vancomycin, render these infections difficult to treat and require prolonged treatment and extended hospitalization ^3-5^.

Additionally, *S. aureus* can survive antibiotic therapy and cause chronic and recurrent infections even while remaining susceptible to antibiotics ^6-8^. This phenomenon of antibiotic persistence or heterotolerance, is defined as the ability of a subpopulation of bacteria to survive high concentrations of bactericidal drugs to which they remain fully susceptible ^9^. Subpopulations with different levels of tolerance can co-exist within a bacterial population, facilitating survival in changing environments ^9,10^. The strategy of bacterial bet hedging via the formation of persister cells in a clonal bacterial population might be advantageous during infection. The actively growing bacteria in a clonal population are mostly responsible for the colonization of the host and a successful infection, while the subpopulation of persisters ensures the survival of the genotype under fluctuating milieus as found in a host and during antibiotic therapy ^11^.

Bacterial persisters can occur spontaneously in a population or can be triggered by environmental factors such as low pH, starvation, oxidative stress and drugs, e.g., by inducing tolerance-by-lag ^12-17^. Signal transduction through reversible protein phosphorylation is a major regulatory mechanism that facilitates quick response to environmental changes ^18^. In bacteria, so called two-component systems, consisting of a histidine kinase sensor and an associated response regulator ^19^, are the most common mechanism of signal transduction. However, regulatory phosphorylation on serine, threonine, and tyrosine residues has been identified in prokaryotes ^20,21^. In *S. aureus*, the conserved eukaryotic-type serine/threonine kinase STK (alternatively named PknB or Stk1) and the cognate phosphatase Stp impacts bacterial cell signalling, central metabolism ^22-24^, stress response ^24,25^, antibiotic resistance ^24,26-28^ and virulence ^26,27,29-31^. Ser/Thr phosphorylation is also involved in the regulation of protein synthesis, the most energy consuming cellular process and both the initiation and elongation phases of translation are downregulated in response to nutrient limitation ^32^. For example, in *B. subtilis*, protein synthesis is downregulated by phosphorylation of the elongation factor Tu (EF-Tu), impairing its essential GTPase activity during nutrient limitation and subsequent formation of dormant spores ^33^.

Here, we describe the role of the Ser/Thr kinase/phosphatase pair PknB/Stp in regulating growth and antibiotic persistence in *S. aureus*. By analysing colony size heterogeneity as well as growth delays of individual bacterial cells using single-cell microscopy, we show that increased serine/threonine phosphorylation leads to the formation of non-stable small colonies (nsSCs) as well as prolonged bacterial lag-phases after acidic pH-stress and in bacteria derived from murine abscesses. The identified growth delay correlates with an increased survival of the bacteria in subsequent antibiotic challenges. Phosphopeptide enrichment via Fe^3+^-Immobilized Metal Affinity Chromatography (IMAC) followed by liquid chromatography-mass spectrometry (LC-MS/MS) analysis enabled us to create a detailed map of PknB and Stp targets. Additionally, we identified changes in phosphorylation states, a decrease in translation activity and increased antibiotic tolerance specifically after acidic pH stress. Together, our results advance the understanding on how *S. aureus* can survive antibiotic treatments despite being fully susceptible, by identifying phosphorylation targets that lead to reduced protein synthesis and cellular quiescence.

## Results

### Deletion of *stp* increases colony growth heterogeneity during stress exposure

*S. aureus* encounters a variety of pH conditions during infection, e.g., a neutral pH of 7.4 in the bloodstream during bacteraemia or an acidic pH on the skin, intracellularly in phagosomes or in abscesses ^34^. We recently demonstrated that so-called non-stable small colonies (nsSCs), whose size is 5 to 10-times smaller than the most common colony type, result from a bacterial growth delay and correlate with the presence of a persister cell subpopulation ^13^. Here, we determined the effect of serine/threonine (Ser/Thr) phosphorylation status on *S. aureus* colony growth and bacterial lag-phases by generating isogenic mutants for *stp* and *pknB* in two distinctive wildtype strains, Cowan I (sequence type [ST] 30) and USA300_JE2 (ST8). These were exposed to defined stress in pH-buffered medium (pH 7.4, 5.5 or 4.0) for three days. While we did not identify any significant differences in the proportion of nsSCs between the wildtype strains and their corresponding isogenic mutants at pH 7.4, we found a significantly elevated proportion of nsSCs in the Δ*stp* mutants (13 to 22%) as compared to the wildtype (5 to 14%) and Δ*pknB* (1 to 15%) after exposure to pH 5.5 for three days (**Fig. 1A**). This effect was even more pronounced when bacteria were exposed to pH of 4.0, with nsSCs levels of up to 37% in the Δ*stp* mutants (**Fig. 1A**). Additionally, there was a trend towards reduced nSCs formation in both Cowan I and USA300_JE2 Δ*pknB* after pH 4 exposure, however not statistically significant (**Fig. 1A**). Furthermore, the deletion mutants Δ*stp* and Δ*pknB* did not exhibit any significant differences in their ability to grow under nutrient rich as well as under various pH conditions when compared to wildtype parent (**Fig. S1, S2A+B**).

**Figure 1:**
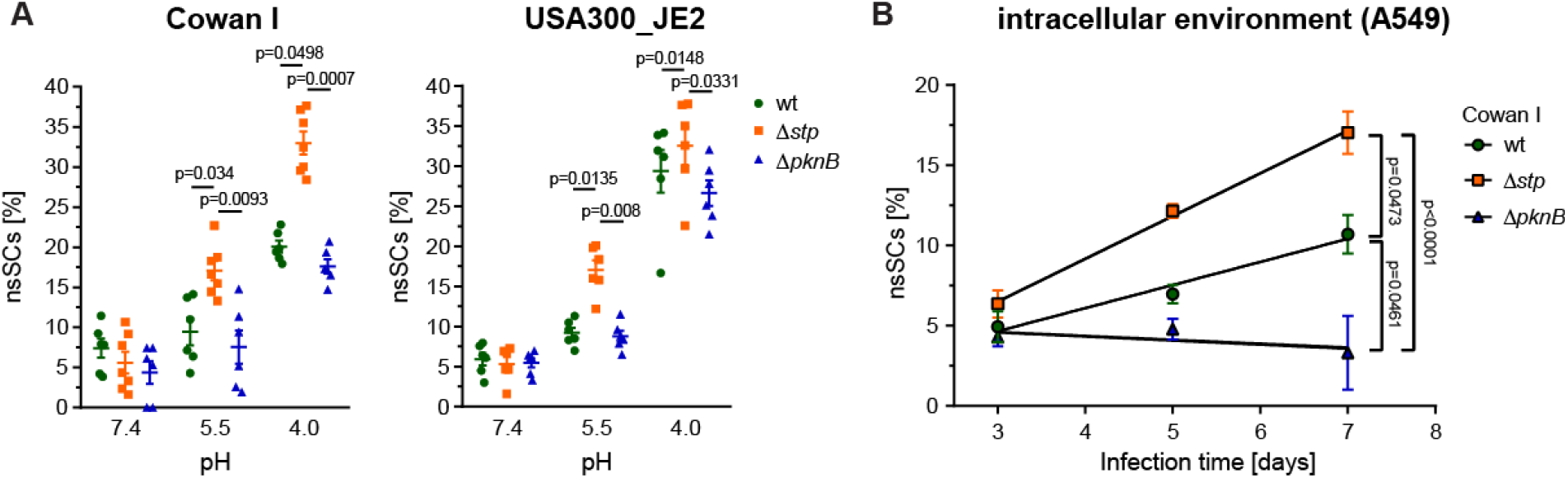
Deletion of *stp* leads to increased formation of nsSCs in *S. aureus* after different stress exposures. **A**) Proportion of nsSCs in *S. aureus* strains Cowan I and USA300_JE2 after three days of pH 7.4, 5.5 or 4.0 exposure. N= 6 to 7 biological replicates. **B)** Proportion of nsSCs in *S. aureus* strain Cowan I after exposure to the intracellular environment of human lung epithelial cells (A549) for 3, 5 and 7 days. Statistically significant p-values are displayed. Statistical significance was determined by the Kruskal-Wallis test with Dunn’s post-test. Error bars depict standard error of mean (SEM).

Exposure to conditions representative of the intracellular milieu, such as acidic pH, can induce the formation of nsSCs with levels increasing over time ^35,36^. To evaluate whether the mutant lacking Stp displays an increased colony size heterogeneity under more complex stress conditions, we infected human lung epithelial cells (A549) with Cowan I and its isogenic mutant strains. Cowan I strain elicited significantly less cytotoxicity towards epithelial cells as compared to the haemolytic USA300 strains, hence was utilized for the intracellular stress assay (**Fig. S3**). We monitored the formation of nsSCs over time 3, 5 and 7 days post infection (p.i.) and found that the Δ*stp* mutant strain formed a significantly higher proportion of nsSCs after 5 and 7 days p.i. as compared to its wildtype or Δ*pknB* counterparts (**Fig. 1B**). In contrast, the kinase deletion mutant Δ*pknB* showed a reduction in nsSCs formation over time as compared to both wildtype and Δ*stp* (**Fig. 1B**). None of the mutants displayed a significantly increased or decreased ability to survive intracellularly, but with a trend towards a reduced capacity to survive intracellularly for Δ*pknB* when compared to the wildtype and Δ*stp* for day 5 and 7 p.i. (**Fig. S2C**). Together, our data suggest that both the phosphatase Stp and the kinase PknB play a role in the *S. aureus* stress response and influence bacterial growth during different stress conditions.

### Growth-resumption delays and reduced ATP-levels in Δ*stp* after acid stress

To verify that the proportion of nsSCs reflects the growth-delay of individual bacterial cells, we determined the time single bacterial cells needed to start their first cell division after changing to favourable growth conditions using time lapse microscopy. The lag-phase was assessed for pH 5.5-exposed bacteria as well as for exponential phase bacteria that served as a control (**Fig. 2A, S4**). While exponentially growing bacteria did not exhibit differences (**Fig. S4**), the lag-phase of mutants after acid stress changed significantly, as expected from our persister measurements on nsSCs (**Fig. 2A**). We observed growth delays of up to 12 h for Cowan I Δ*stp* and 10 h for USA300_JE2 Δ*stp*. The wildtype strains showed lag-phases of up to 8 h for Cowan I and 4 h for USA300_JE2. While there was no significant difference in the lag-phases between the wildtype and Δ*pknB* in Cowan I (90% of the bacteria have started to grow after 5.5 h), USA300_JE2 Δ*pknB* had a significantly reduced lag-phase as compared to its wildtype counterpart after pH 5.5 exposure (**Fig. 2A**).

**Figure 2:**
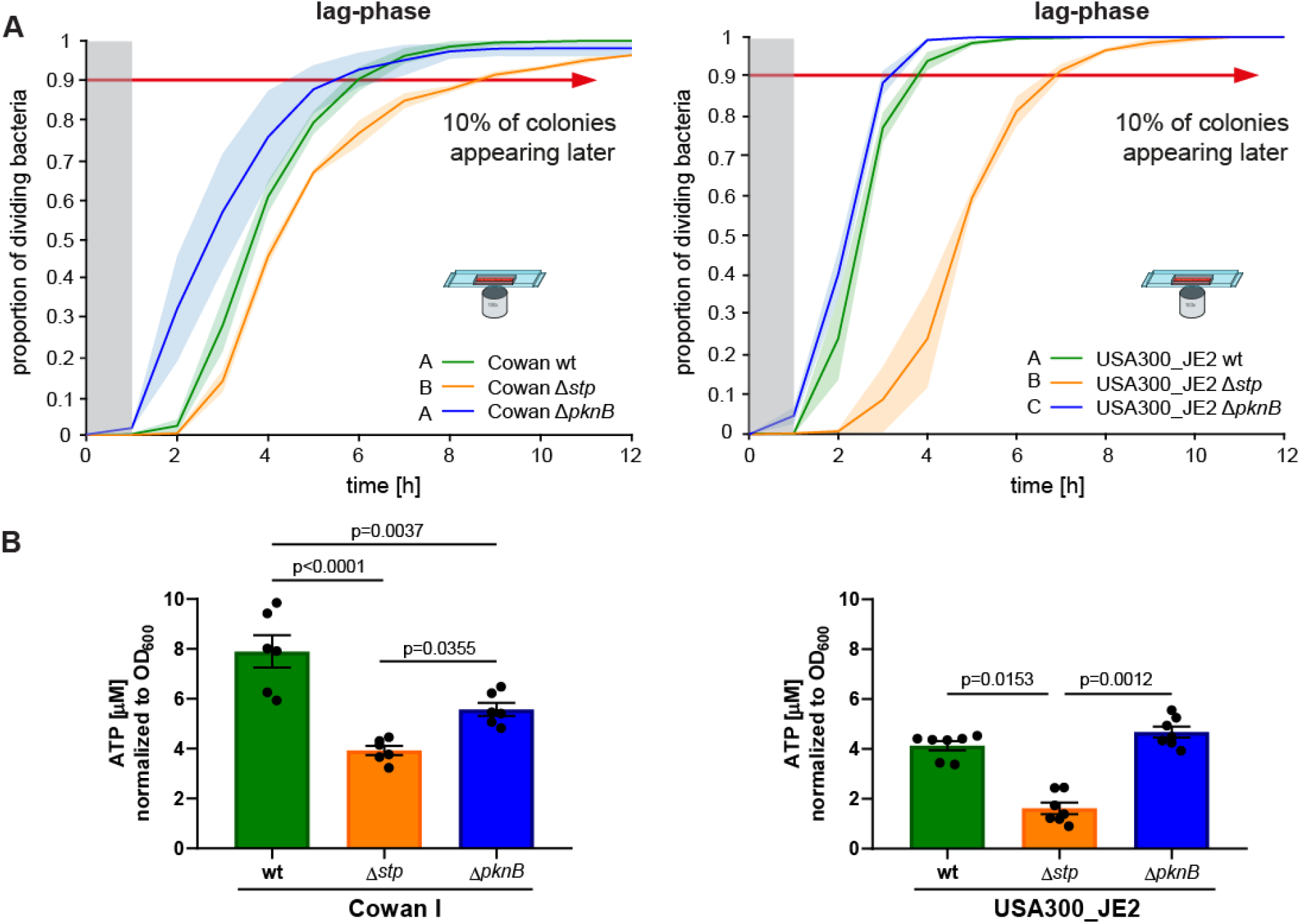
Acidic pH-exposure increases bacterial lag-phase and results in reduced ATP levels in *S. aureus* Δ*stp*. **A**) Single-cell microscopy of *S. aureus* wildtype and the isogenic mutants Δ*stp* and Δ*pknB* growing in nutrient-rich medium after three days of pH 5.5 exposure. N= 3 biological replicates. Shaded areas depict the standard deviation. Cowan I wt pH 5.5 n= 475 bacterial cells, Cowan I Δ*stp* pH 5.5 n= 246, Cowan I Δ*pknB* pH 5.5 n= 554; USA300_JE2 wt pH 5.5 n= 662 bacterial cells, USA300_JE2 Δ*stp* pH 5.5 n= 669, USA300_JE2 Δ*pknB* pH 5.5 n= 1,062; Grey zone marks the period at the beginning of the experiment, where cell divisions could occur, but would not be observed. Letters indicate statistically significantly different groups. Statistical significance was determined for the time points where 90% of the bacteria started to divide (red arrow). **B**) *S. aureus* was grown in DMEM pH 5.5 for three days. Intracellular ATP concentration was determined using a BacTiter-Glo cell viability assay. N= 6 biological replicates. Statistically significant p-values are displayed; Statistical significance was determined by one-way ANOVA with Tukey post-test after passing the Shapiro-Wilk normality test. Error bars depict SEM.

To determine whether the growth delay is linked to reduced energy levels, we measured intracellular ATP concentrations of the bacteria after exposure to acidic pH (**Fig. 2B**). We found that the Δ*stp* mutant, which showed significantly prolonged bacterial lag-phases after pH 5.5-exposure (**Fig. 2A**), displayed significantly lower ATP levels (50% reduction) as compared to the wildtype and Δ*pknB* mutant in both Cowan I and USA300_JE2 (**Fig. 2B**). Slightly, but significantly reduced ATP levels were found for Cowan Δ*pknB* as compared to the wildtype, while there was no difference between USA300_JE2 wildtype as compared to Δ*pknB*. These findings are consistent with previously published metabolomics data showing significantly reduced ATP levels in stationary *S. aureus* strain 8325 Δ*stp*, but not in Δ*pknB* ^37^. Our data show that, in the absence of Stp, the single cell growth delay after acid stress is associated with reduced intracellular ATP levels.

### Delays in growth resumption and reduced ATP levels contribute to antibiotic persistence

To determine if high proportions of late-growing bacteria with reduced ATP levels are associated with increased antibiotic persistence, we challenged acidic pH-exposed wildtype strains of *S. aureus* as well as their deletion mutants Δ*stp* and Δ*pknB* with high concentrations of antibiotics (40x the minimum inhibitory concentration [MIC]). To test the survival capacity of Cowan I, we used the cell wall targeting antibiotic flucloxacillin (FLU), the DNA-replication inhibitor fluoroquinolone ciprofloxacin (CIP) and the transcription inhibitor rifampicin (RIF). For USA300_JE2, we used the cell wall targeting drugs FLU and vancomycin (VAN) as well as RIF.

Consistent with the increased bacterial lag-phase and the reduced ATP levels in the Δ*stp* mutants of both strains after pH 5.5-exposure (see **Fig. 1**+**2**), we observed increased levels of antibiotic tolerance towards several different antibiotics (**Fig. 3A-D**). Calculating the minimum duration for killing 90% of the initial bacterial population (MDK_90_) for FLU and CIP as well as the MDK_99_ for the rapidly acting antibiotic RIF for Cowan I (**Fig. 3B**) and MDK_90_ for FLU, VAN and RIF for USA300_JE2 (**Fig. 3D**) allowed us to better compare between strains and treatments. We found that 90% of the initial population of pH 5.5-exposed Cowan Δ*stp* was killed by FLU after 19 h, whereas it took only 6 h for the wildtype and 5 h for Δ*pknB*. A similar pattern was revealed for CIP and RIF (**Fig. 3B**). Analogously, a similar behaviour was identified for USA300_JE2 challenged with FLU, VAN or RIF (**Fig. 3D**). To exclude resistance development as a confounding factor for RIF persistence, we additionally plated on RIF-plates and excluded experiments where resistance development occurred from this analysis (**Fig. S5**).

**Figure 3:**
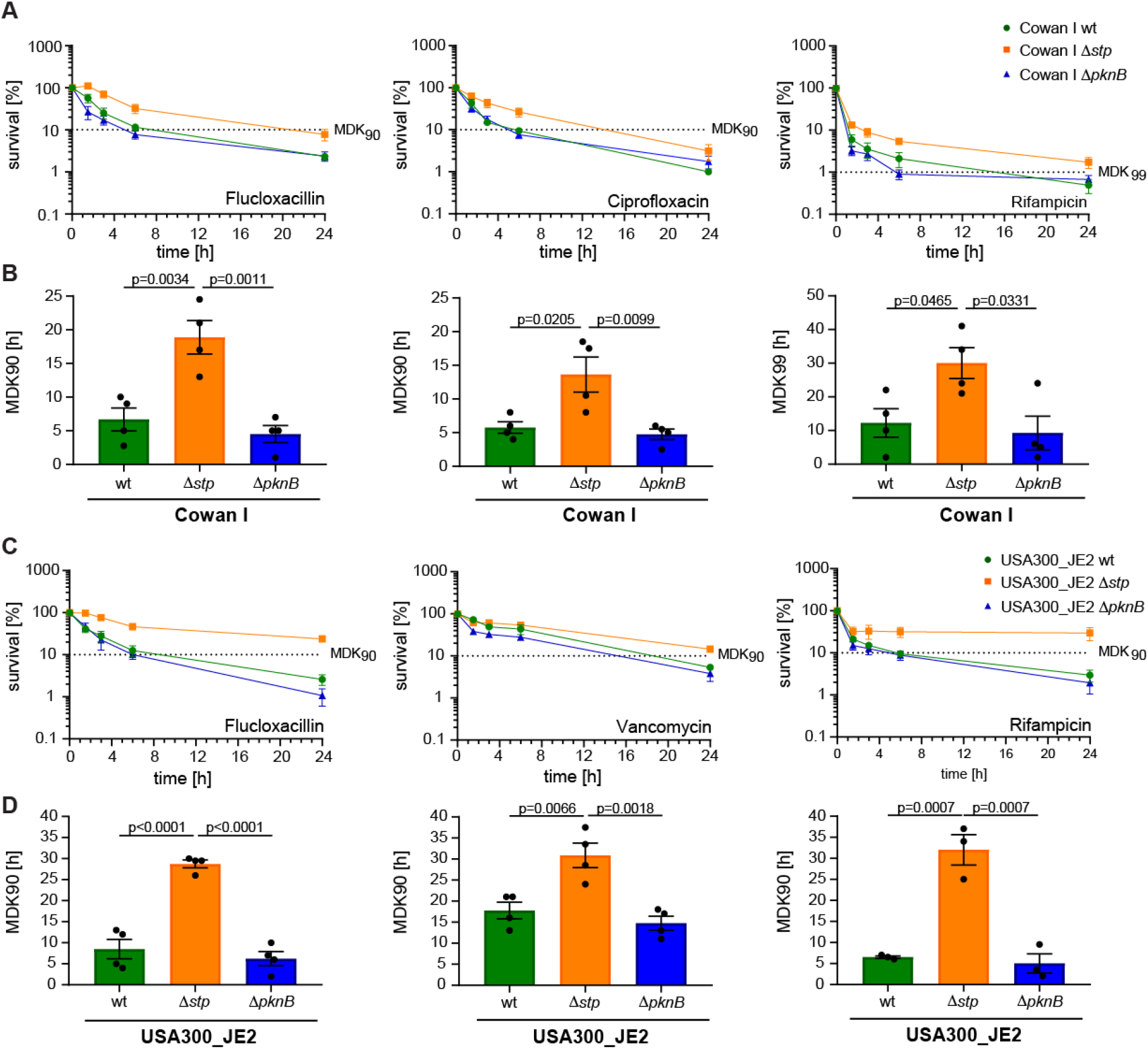
Increased antibiotic tolerance in Δ*stp* after acidic pH-exposure. **A-D**) *S. aureus* was grown in DMEM pH 5.5 for three days before inoculation in neutral pH medium supplemented with antibiotics (40xMIC) as indicated. **A+C**) Time kill curves (0, 1.5, 3, 6 and 24 h) for different antibiotics and strains (Cowan I and USA300_JE2). To determine bacterial survival, bacteria were sampled at different time points, washed three times to eliminate antibiotics and plated for CFU enumeration. Survival was calculated relative to the initial inoculum of ∼10^5^ CFUs/ml. **B+D**) Minimum duration of killing 90 or 99% of the initial population for different strains and antibiotics. Statistically significant p-values are depicted. Statistical significance was determined by one-way ANOVA with Tukey’s multiple comparison testing after passing the Shapiro-Wilk normality test. Error bars depict SEM.

### *stp* deletion leads to increased bacterial lag-phase and antibiotic persistence in mice

To test whether the *stp* deletion leads to increased bacterial lag-phases and a subpopulation of persisters *in vivo*, we infected mice with either *S. aureus* wildtype or the corresponding isogenic mutants Δ*stp* and Δ*pknB* (**Fig. 4A**). Here, we used the less cytotoxic Cowan I to induce abscess formation and pus production in the mice as described previously ^13,38^. Five days post-infection, mice were sacrificed, abscess size measured, and the pus harvested for further analysis. The Δ*stp* strain caused significantly larger abscesses compared to Δ*pknB*, but there was no significant difference between the wildtype and Δ*stp* as well as wildtype and Δ*pknB* (**Fig. S6A**). Analysis of the colony growth heterogeneity showed that Δ*stp* formed a higher proportion of nsSCs (up to 13%) as compared to both wildtype (up to 5.9%) and Δ*pknB* (up to 4.8%), indicating the presence of a larger quiescent subpopulation in the absence of Stp in *in vivo* abscess settings (**Fig. 4B, S6B+C**). Single cell microscopy confirmed prolonged individual bacterial lag-phases in Δ*stp* as reflected by the increased proportion of nsSCs (**Fig. 4C**). To test whether *S. aureus* lacking the phosphatase Stp not only forms larger abscesses and more nsSCs, but also a larger subpopulation of persister cells, we exposed bacteria isolated from murine abscesses to high concentrations of FLU and CIP (40xMIC) for 24 h and analysed their survival capacity (**Fig. 4D**). We found that the Δ*stp* mutant showed significantly higher survival levels (up to 16%) as compared to the wildtype (up to 6%), correlating with the increased proportion of nsSCs (**Fig. 4E, S6D+E)**. Additionally, the kinase mutant Δ*pknB* showed reduced antibiotic persistence; we found significantly lower survival in FLU as compared to both wildtype and Δ*stp* as well as a trend to lower survival compared to the wildtype in CIP (**Fig. 4D**). Together, our data demonstrate that the kinase/phosphatase pair PknB/Stp modulates antibiotic persistence in *S. aureus in vivo*, and that a lack of Ser/Thr dephosphorylation can lead to an increased subpopulation of persisters.

**Figure 4:**
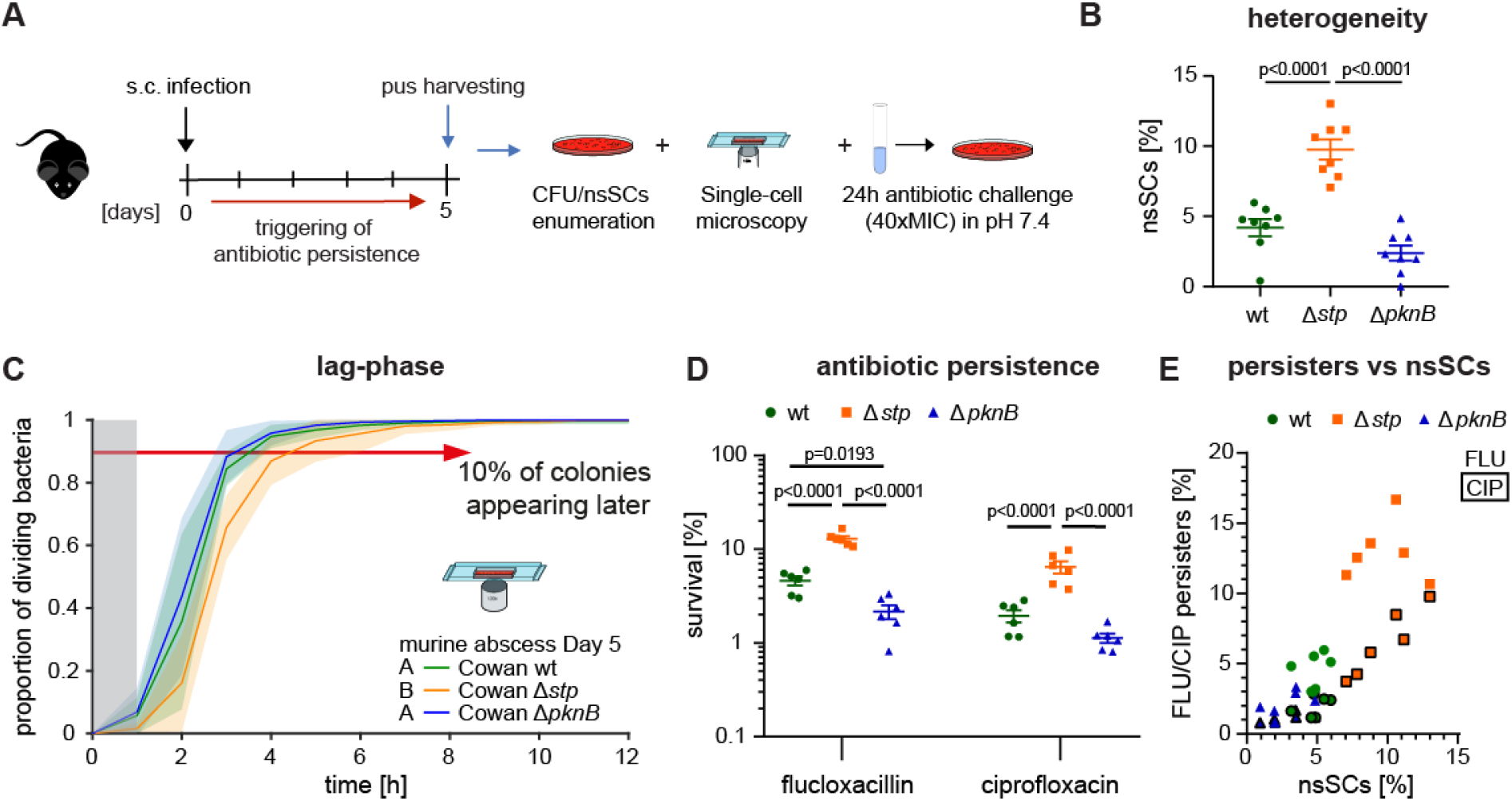
Deletion of *stp* leads to increased bacterial lag-phase and antibiotic persistence in mice. **A**) Scheme of the murine infection model. **B**) Colony size heterogeneity in murine abscess pus. Proportion of nsSCs was calculated relative to total CFU (**Fig. S6B+C**). Statistical significance was determined by one-way ANOVA with Tukey’s multiple comparison testing. **C**) Single cell microscopy to determine individual cells’ lag-phases after recovery from murine abscess pus. Cowan I wildtype n= 727 bacterial cells; Cowan I Δ*stp* n= 1,038 and Cowan I Δ*pknB* n= 776; Grey zone marks the period at the beginning of the experiment where cell divisions could occur but would not be observed. Letters indicate statistically significantly different groups. **D**) Survival assay of bacteria harvested from murine abscesses and exposed to FLU or CIP for 24 h. Survival was calculated relative to the initial inoculum. Statistical significance was determined by a two-way ANOVA with Tukey’
ss multiple comparison testing after passing the Shapiro-Wilk normality test. **E)** Relationship between the proportion of nsSCs and the proportion of persisters (FLU= no border, and CIP= black boarder colour). Each symbol represents one murine abscess. **B-E)** N= 6 abscesses from 3 mice. Error bars depict SEM.

### Phosphoproteomic analysis reveals protein synthesis as a target of PknB and Stp during acid stress

To elucidate the role of Ser/Thr phosphorylation in *S. aureus* antibiotic persistence and growth regulation, and to further identify potential down-stream targets of PknB and Stp, we performed total proteome as well as phosphoproteomic analyses on pH 5.5 exposed strain Cowan I wt and its respective *pknB* and *stp* deletion mutants (**Fig. 5A**). As previously reported ^39^, *S. aureus* favours single phosphorylated peptides, with over 80% of identified peptides phosphorylated only once. However, deletion of Stp led to an increased number of single and double phosphorylated peptides, while deletion of *pknB* showed no drastic differences when compared to the wildtype (**Fig. S7A**), suggesting the presence of additional Ser or Thr kinases in *S. aureus*. Similarly, a heatmap overview of all identified phosphopeptides (**Fig. 5B**) showed a similar pattern in wildtype and Δ*pknB*, with only 33 significantly underrepresented and seven overrepresented phosphopeptides in Δ*pknB* as compared to wildtype (**Fig. S7B**, *right panel*, and **D**). A prominent difference in phosphopeptides was revealed between Δ*stp* and wildtype, with 240 phosphopeptides significantly overrepresented (90% on Thr) and 137 significantly underrepresented (55.5% on Ser) (**Fig. 5C**, *right panel*, and **S7B**). On the entire proteome level, 1,550 proteins (coverage of 59.5%) were identified, of which 2.8% (43 proteins) were differentially expressed. Specifically, 1.5% (23 proteins) were found to be significantly downregulated and 1.3% (20 proteins) significantly upregulated in Δ*stp* as compared to wildtype (**Fig. 5C**, *left panel*). Consequently, most of the changes occurred exclusively on the phosphoproteome and not on the total proteome (**Fig. S7C**), which is in line with previously published data ^39^. Phosphorylation of ribosomal proteins such as RplS (pT11), RpsC (pT176) and RpsP (pT70), translation elongation factors EF-Tu (pT34) and EF-G (pT34, pT43 and pT424) and cell division proteins such as FtsZ (pT333, pS338 and pT367) were prominently overrepresented in the *stp* deletion mutant as compared to the wildtype (**Fig. 5C**, *right panel*, and **5D+E**). Deletion of the kinase PknB, on the other hand, showed a similar phosphorylation pattern as the wildtype for both ribosomal proteins and the translation elongation factor EF-G (**Fig. S7E**+**F**). Phosphorylation of translation elongation factors has been linked to reduced translational activity and bacterial dormancy in *B. subtilis* ^33^ and our data shows that phosphorylation of translation elongation factors seems to play an important role in *S. aureus* as well (**Fig. 5**). The increased phosphorylation on several Thr-residues in EF-G, e.g., the sixteen-fold increase for pT43, was confirmed by immunoblotting wherein only in Δ*stp* the antibody signal from EF-G and P-Thr overlapped but did not align in the wildtype nor in Δ*pknB* (**Fig. 5F, S7G**). Similar results were found for *S. aureus* USA300_JE2 (**Fig. S7H**).

**Fig. 5:**
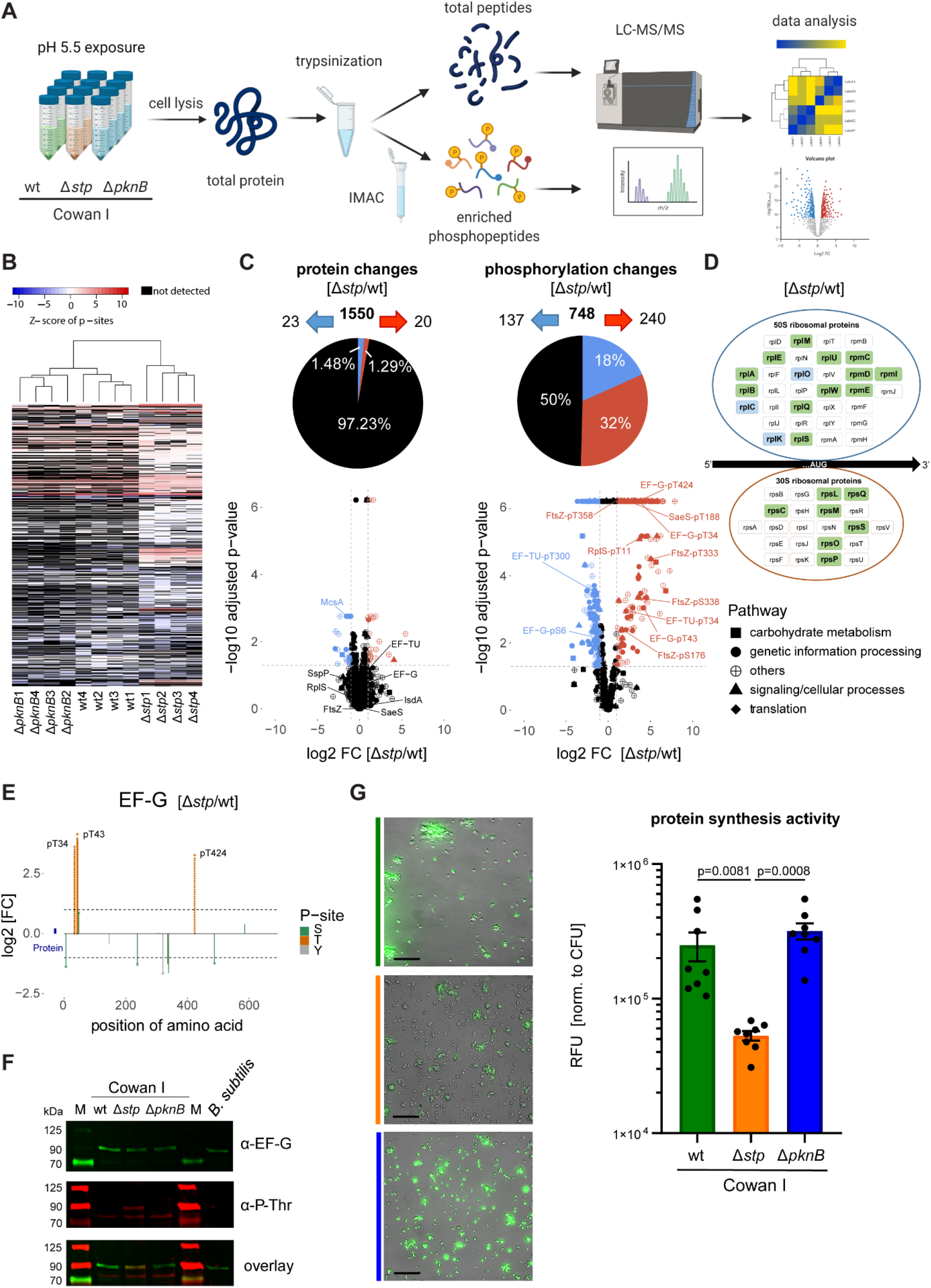
Phosphorylation and protein expression alterations in acid stress exposed *S. aureus*. **A)** Experimental workflow for proteomic and phosphoproteomic analysis. After cell lysis to isolate bacterial total protein from *S. aureus* strain Cowan I grown at pH 5.5 for three days, total proteome and enriched phospho-peptides were analysed via label-free LC-MS/MS. **B)** Hierarchical cluster analysis of phosphopeptide abundance in phosphoproteome. The abundance profiles of the phosphopeptides in our four replicates hierarchically clustered by heatmap. Blue indicates less abundant phosphopeptides, whereas red indicates more abundant phosphopeptides. Black indicates non-detected phosphopeptides. **C)** Pie charts and volcano plots comparing total proteome (*left panel*) and phosphopeptide (*right panel*) changes between Cowan I *stp* deletion mutant and wildtype, identifying differentially expressed proteins or phosphopeptides (cut-off adjusted p-value < 0.05 and a fold change of |log_2_FC| > 1). Underrepresented proteins/ phosphopeptides are depicted by blue symbols and overrepresented proteins/ phosphopeptides with red symbols. Symbol type indicates Kyoto Encyclopedia of Genes and Genomes (KEGG) pathway. **D)** Identified changes in Ser/Thr phosphorylation of ribosomal proteins in Cowan I *stp* deletion mutant compared to its wildtype. Green colour indicates increased Ser/Thr phosphorylation (at least one phospho-site log_2_FC ≧ 1, adjusted p-value < 0.05). Blue colour indicates decreased Ser/Thr phosphorylation (at least one phospho-site log_2_FC ≦ -1, adjusted p-value < 0.05). If upregulated and downregulated phosphopeptides for the same protein were found, the highest up- or downregulation is shown. **E)** P-sites to protein plot for the elongation factor EF-G. Phosphorylation sites and exact amino acid positions are indicated. Y-axis indicates log_2_ fold changes of phosphopeptides in Cowan I Δ*stp* compared to wildtype. Dashed lines indicate pseudo fold changes (phosphopeptides were only detected in one condition). **F)** Representative immunoblots of bacterial cell lysates with antibody specific for phospho-threonine and rabbit serum raised against *B. subtilis* EF-G that also recognises *S. aureus* EF-G. Cowan I wildtype, Δ*stp* and Δ*pknB* were grown at pH 5.5 for three days, *B. subtilis* was grown to stationary phase for 16 h. **G)** Representative images of OPP-stained Cowan I wildtype, Δ*stp* and Δ*pknB*. Fluorescence indicates protein synthesis (*left panel*). Total fluorescence of OPP-labelled *S. aureus* normalized to CFUs (*right panel*). Scale bar indicates 20 µm.

To assess if the overrepresented phosphorylation of ribosomal and translation elongation factors is associated with differences in protein synthesis activity, we measured incorporation of the puromycin analogue O-propargyl-puromycin (OPP) ^40^ to monitor protein synthesis in pH 5.5 exposed *S. aureus* Cowan I. We found that the *stp* deletion mutant shows significantly reduced protein synthesis activity as compared to the wildtype, while the *pknB* deletion mutant displayed increased protein synthesis (**Fig. 5G**). Taken together, these data suggest that phosphoregulation of proteins involved in translation leads to reduced protein synthesis in *S. aureus* during acid stress exposure.

## Discussion

This study shows that the Ser/Thr kinase PknB and its cognate Ser/Thr phosphatase Stp regulates antibiotic persistence in *S. aureus* (as summarized in **Fig. 6)**. Measurement of translation showed significantly reduced protein synthesis consistent with increased Ser/Thr phosphorylation after acid stress when the phosphatase Stp is absent. Similarly, deletion of *stp* increases the subpopulation of quiescent bacteria upon stress exposure, which was reflected by elevated levels of nsSCs and by lag-phase analysis of individual bacterial cells using single cell microscopy. Additional analysis revealed decreased levels of intracellular ATP in the Δ*stp* mutant, further indicating a metabolically reduced state of the bacteria. Both *in vitro* acid stress, and *in vivo* stress exposure in murine abscesses lead to increased numbers of bacteria with a high growth delay. Our data show that the growth delay as well as the reduced ATP levels of an Δ*stp* mutant relates to its increased ability to survive high concentrations of antibiotics from different classes. Finally, phosphoproteomic analysis of the *S. aureus* strain Cowan I revealed increased phosphorylation of proteins involved in translation such as ribosomal proteins and translation factors. This highlights the role of phospho-regulation in bacterial quiescence and survival during antibiotic challenges and the interplay between the kinase PknB and the phosphatase Stp (**Fig. 6, S10)**.

**Figure 6:**
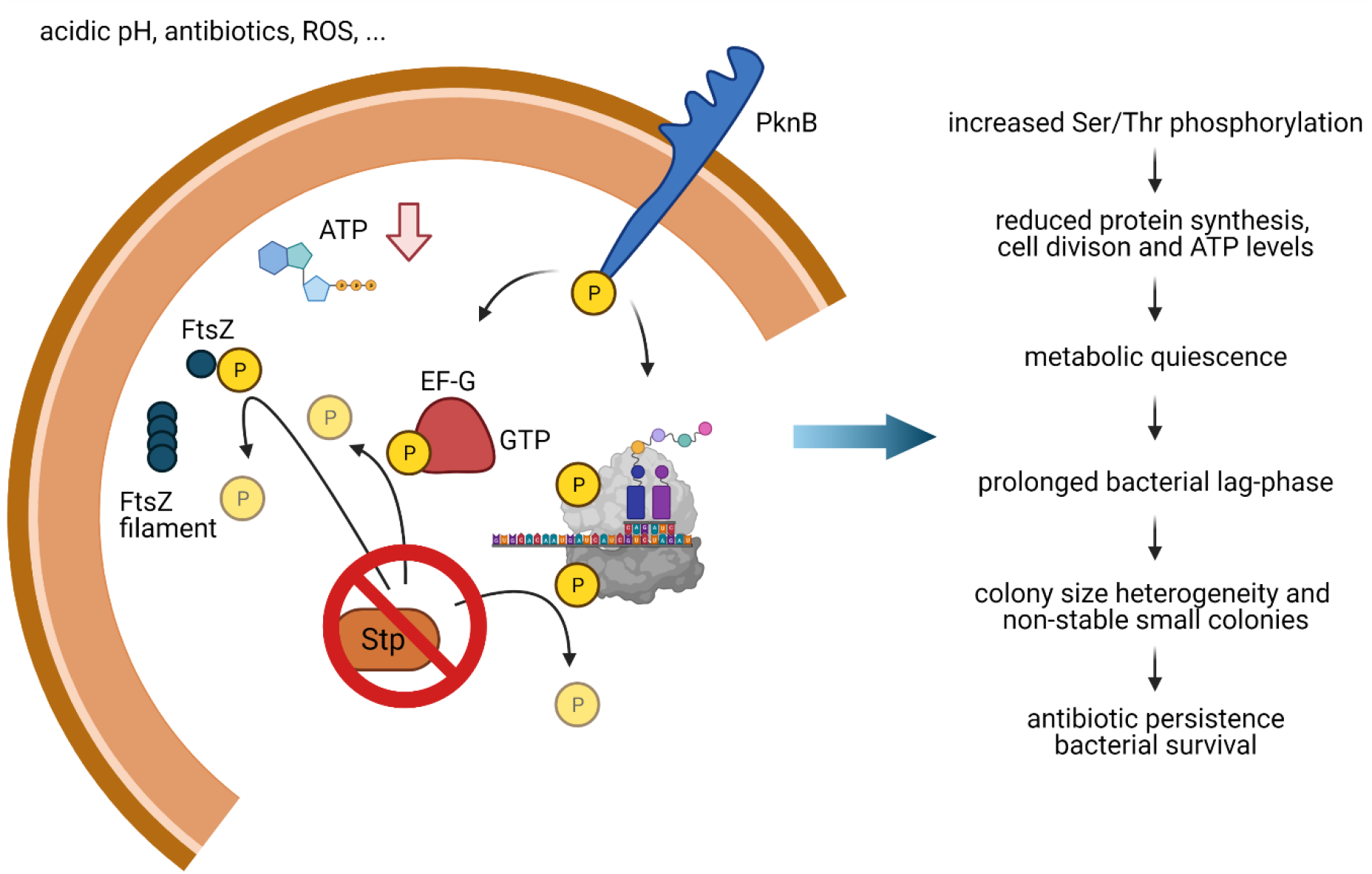
Phosphorylation-changes linked to cellular quiescence and antibiotic persistence in *S. aureus*. Scheme summarizing the results of this study (see **Fig. S10**). Sensing of extracellular stressors by the eukaryote-like Ser/Thr kinase PknB leads to phosphorylation of proteins involved in protein synthesis, cell division and growth in *S. aureus* resulting in reduced protein synthesis, cell division and ATP levels. This metabolic quiescence leads to prolonged bacterial lag-phases and thus to a delayed onset of growth after switching to favourable conditions. This growth delay manifests in the formation of non-stable small colonies (nsSCs) and eventually in colony size heterogeneity on agar plates. Ultimately, *S. aureus* is able to survive antibiotic challenges and shows increased antibiotic persistence. Yellow circles indicate phosphor residues. ATP= adenosine triphosphate, Stp= Ser/Thr phosphatase, EF-G= elongation factor G, FtsZ= core divisome component protein, GTP= guanosine triphosphate; Created with BioRender.com.

PknB was initially identified in a genetic screen designed to detect factors that regulate methicillin resistance in *S. aureus* ^41^. It is highly conserved throughout all sequenced *S. aureus* genomes ^22,30,42^ and has a domain architecture consisting of extracellular penicillin-binding protein- and STK-associated (PASTA) domains, a transmembrane domain and an intracellular catalytic kinase domain, similar to *B. subtilis* PrkC as well as *M. tuberculosis* PknB ^31^. Previous phosphoproteomic studies identified PknB target proteins involved in metabolic processes such as glycolysis, citrate cycle, cell wall synthesis and protein synthesis ^23,39,43^. Both PknB and Stp play an important role in cell wall synthesis and it has been shown that deletion of *stp* leads to a thickened and more robust cell wall in *S. aureus* resulting in decreased susceptibility to lysostaphin and vancomycin ^27,42-44^. Similarly, we observed a two-fold increase in the MICs for cell wall targeting antibiotics such as flucloxacillin and vancomycin (**Table S3**). However, as we used high antibiotic concentrations for our killing assays, doubled MIC values for ß-lactams do not alone explain the increased survival of the *stp* deletion mutants during ß-lactam challenge.

We recently found elevated levels of proteins involved in protein, nucleotide and amino-acid biosynthesis in a persister-enriched population of a clinical isolate of *S. aureus* that was isolated from a difficult-to-treat infection, ^38^. Higher levels of PknB might lead to increased phosphorylation levels that in turn can reduce translation, alter cell wall synthesis and can lead to decreased intracellular ATP levels. Inhibition of protein synthesis by phosphorylation of the elongation factor Tu (EF-Tu) in *B. subtilis* has been shown to occur during nutrient limitation and ultimately lead to the formation of dormant spores-an extreme form of bacterial persistence ^33^. In *E. coli*, phosphorylation of EF-Tu leads to its inactivation and to the inhibition of protein synthesis during environmental stress such as starvation ^45^. In general, inhibition of protein synthesis has been shown to increase antibiotic persistence in *E. coli* ^46,47^.

Here, we show that deletion of *stp* leads to increased Thr phosphorylation (pT34, pT43, pT424) of EF-G and extensive phosphorylation of ribosomal proteins in acid stress exposed *S. aureus*. Together with reduced protein synthesis, low ATP levels and a delayed onset of growth after stress removal, this correlated with increased antibiotic tolerance in *S. aureus*. Also, proteins involved in cell division and bacterial growth were found to be phosphorylated by PknB. For example, the core divisome component FtsZ showed increased phosphorylation in the absence of Stp after acid-stress exposure. Phosphorylation of FtsZ might inhibit its GTPase activity leading to arrested cell division ^48,49^. Furthermore, it has been shown that Ser/Thr phosphorylation of FtsZ is important for complex formation with the cell division protein FtsQ in *M. tuberculosis* during oxidative stress, highlighting the importance of Ser/Thr phosphorylation in maintaining bacteria in a division-competent state depending on the growth conditions ^50^.

Bacterial growth delays and cellular quiescence have been linked to antibiotic tolerance and the presence of nsSCs can be indicative of a subpopulation of persister cells ^13,51,52^. The increased levels of nsSCs in *S. aureus* lacking Stp and the growth delay observed on the single-cell level are associated with reduced energy levels and increased bacterial survival during antibiotic exposure. Reduced intracellular ATP levels have been linked to antibiotic tolerance as most antibiotics require an active target and bacterial metabolism in order to eradicate bacteria ^53-55^. Growth delays, nsSCs as well as antibiotic persistence were not only detected after *in vitro* stress exposure, but also *in vivo* after harvesting bacteria from murine abscesses. Since deletion of the kinase PknB did not completely eliminate Ser/Thr phosphorylation in *S. aureus*, other phospho-donors such as acetyl-phosphate, ammonium hydrogen and carbamoyl phosphate or other currently unknown Ser/Thr kinases ^39,56,57^ may participate in this response.

We generated a comprehensive map of the phosphoproteome of *S. aureus* Cowan I and its isogenic mutant strains exposed to acidic pH linking Ser/Thr phosphorylation to reduced protein synthesis and intracellular ATP levels and thereby increased bacterial survival during antibiotic exposure. The kinase inhibitor staurosporine increases methicillin-resistant *S. aureus* (MRSA) sensitivity to the β-lactam nafcillin ^58^ suggesting that inhibition of PknB during infection might reduce formation of persister cells. Thus, the kinase/phosphatase pair PknB and Stp may be a future target to prevent persister formation during chronic and difficult-to-treat infections that require prolonged antibiotic therapies.

## Materials and Methods

### Bacterial Culture and Growth Conditions

Bacterial strains used in this study are listed in **Table S1**. *S. aureus* was grown on Columbia agar plates supplemented with 5% sheep blood (BioMérieux) overnight prior to use or on tryptic soy broth (TSB) (Becton Dickinson) containing 1.5% agar (Becton Dickinson) for bacterial enumeration. For exponential growth conditions, bacteria were grown in TSB overnight (16 h) shaking (220 rpm) at 37°C, diluted and regrown in TSB, shaking at 37°C, for two hours. For pH-stress assays, *S. aureus* was inoculated into defined pH media (Dulbecco’s Modified Eagle Medium (DMEM) pH 4.0, 5.5 and 7.4) at a starting OD_600_ of 0.2. After incubation at 37°C and 5% CO2 for three days and daily mixing, cultures were analysed by plating and counting nsSCs or by determining antibiotic persistence in a survival assay. NsSCs enumeration was performed as described previously ^13^. Specific growth conditions are described in the corresponding method sections.

### Genetic manipulations

For both strains, Cowan I and USA300_JE2, unmarked deletions for either *pknB* or *stp* were created using the pIMAY vector as previously described ^59^. Briefly, *S. aureus* strains were transformed with pIMAY containing the modified genomic sequences missing either *pknB* (pSP11; **Fig. S8**) or *stp* (pSP13; **Fig. S9**) by electroporation. Primers and plasmids used to create pSP11 and pSP13 are described in **Table S2**. Colonies grown on tryptic soy broth agar plates supplemented with 10 µg/ml chloramphenicol (Cm), containing the plasmid were used for chromosomal integration by homologous recombination. Deletion of *pknB* and *stp* was confirmed by PCR and sequencing. To rule-out inherent growth defects of the deletion mutants, we performed growth curves and did not identify any differences in growth of the mutants compared to their wildtypes (**Fig. S1**).

### Eukaryotic cell infections

The human lung epithelial carcinoma cell line A549 (ATCC CCL-185) was grown in DMEM 4.5 g ml^−1^ glucose (Life Technologies) with 10% foetal bovine serum (FBS; GE-Healthcare) and 2 mM L-glutamine (Life Technologies) and infected with *S. aureus* strain Cowan I at a multiplicity of infection of 1. Three hours after infection, the cells were washed with phosphate-buffered saline (DPBS; Life Technologies), and flucloxacillin (Actavis; 1 mg ml^−1^ in DMEM) and lysostaphin (Sigma; 25 µg ml^-1^ in DMEM) were added to kill any extracellular bacteria. Every day, the washing step was repeated, and fresh medium (containing flucloxacillin 1 mg ml^−1^ and lysostaphin 25 µg ml^-1^) was added to the host cells. Supernatants were monitored for the absence of any extracellular bacteria by plating. Three-, five- and seven-days post-infection, host cells were washed three times with PBS, lysed (0.08% Triton X-100 (Fluka) in PBS), and plated on Columbia agar plates supplemented with 5% sheep blood for nsSCs enumeration.

### Single-cell time-lapse microscopy

Agar pads were prepared using a custom-made device made of polyvinyl chloride. Bacterial dilutions were added onto solidified 2% agar (bacteriological grade, BD) with Columbia media (BD) and 5% sheep blood (Life Technologies) and then covered with a cover glass. The setup was then placed under a microscope at 37°C at the latest 30 min after inoculation. Bright field images were acquired using a fully automated Olympus IX81 inverted microscope at 100× resolution (U-FLN-Oil lens) and the Cellsense software. Up to 3,000 positions were monitored per experiments and the lag-phase of 246–2,000 bacteria per strain was analysed ^13^. Statistical significance was determined for time points where 90% of bacteria started to divide using unpaired t-tests.

### Mouse experiment

The protocols (ZH251/14 and ZH050/18) were approved by the institutional animal care and use committee of the University of Zurich and all experiments were conducted in approval of the Cantonal Veterinary Office Zurich. *S. aureus* Cowan I wildtype as well as the isogenic mutants Δ*pknB* and Δ*stp* were grown to logarithmic phase and 10^8^ CFUs mixed 1:1 with cytodex beads (Sigma) ^60^. Bacteria were injected into the flanks of 7–9-weeks-old female C57BL/6 mice (Janvier Laboratory, France). Mice were sacrificed 5 days post-infection. Abscess area was measured, and abscess volume calculated as previously described ^61^. Abscess pus was harvested, homogenized, lysed and directly analysed by single-cell time-lapse microscopy as well as plating serial dilutions for bacterial enumeration and nsSC quantification. Additionally, bacteria from abscess pus were used for subsequent persister assays.

### ATP-assay

Intracellular adenosine triphosphate (ATP) levels were measured using a BacTiter-Glo kit (Promega) according to the manufacturer’s instructions. *S. aureus* grown in DMEM supplemented with 10% FBS and 1% L-Glu at pH 5.5 for three days were washed with PBS before measuring luminescence on a SpectraMax i3 MiniMax 300 plate reader (Molecular Devices) in white 96-well flat bottom plates. (Greiner Bio-One). ATP levels were normalized to CFU counts. N= 6 biological replicates.

### Persister assay and time kill curves

Approximately 10^5^ bacteria from different stress conditions (pH 5.5 or murine abscess pus) were inoculated in DMEM pH 7.4 supplemented with antibiotics (40xMIC) as indicated. After 1.5 h, 3 h, 6 h and 24 h of incubation at 37°C and 5% CO_2_, bacterial survival was determined by CFU enumeration and calculated relative to the inoculum. MICs for FLU, VAN, RIF and CIP were assessed by micro broth dilution method, following the “European Committee for Antimicrobial Susceptibility Testing” (EUCAST) recommendations. Minimum duration of killing 90% or 99% of the initial population (MDK90/99) was calculated for each replicate.

### OPP-staining and translational activity measurement

Click-iT Plus OPP Alexa Fluor™ 488 Protein Synthesis Kit (Invitrogen) was used to label *S. aureus* with O-propargyl-puromycin (OPP) following the manufacturer’s instructions. Briefly, 450 µl of bacteria exposed to pH 5.5 for 3 days were washed with PBS and resuspended in pre-warmed TSB supplemented with 20 µM OPP. Labelling was performed at 37°C and 220 rpm shaking for 20 min. All subsequent steps were performed at room temperature. After incubation, bacteria were washed with PBS and fixed in 3.7% formalin for 15 min. To permeabilize the bacteria, fixative was removed, and bacteria were incubated in 100 µl of 0.5% Triton X-100 in PBS for 15 min. After washing twice with PBS, bacteria were labelled using 1X Click-iT cocktail for 30 min in the dark. Bacteria were harvested and washed once with Click-iT rinse buffer and resuspended in 40 µl of PBS for imaging or 150 µl of PBS for fluorescence measurement on a SpectraMax i3 MiniMax 300 plate reader in white 96-well flat bottom plates.

### LC-MS/MS

#### Proteomics sample preparation

A total of 12 samples (4 biological replicates for each strain) were processed for mass spectrometry-based proteomics analysis. To extract the proteins, 30 µl SDS-lysis buffer [4% (w/v) SDS, 100mM Tris-HCl pH 8.2, 0.1M dithiothreitol complemented with Halt Protease and Phosphatase Inhibitor (Thermo Fisher Scientific)] was added to the frozen cell pellets. Samples were mechanically lysed with glass beads in a tissue lyser (QIAGEN) for 4 cycles of 2 min, running at 30 Hz. Between the cycles, samples were cooled down on ice. After spinning down the samples, samples were boiled for 5 min at 95°C and placed 1 min in high-intensity focused ultrasound sonication at 100% amplitude, 80% cycle (UP200St with VialTweeter, Hielscher Ultrasound Technology). Samples were then diluted with 200 µl 8 M urea, 100 mM Tris-HCl (pH 8.2) and centrifugated for 10 min at 8,000g at 4°C. The clear supernatant was transferred into a new tube. Protein concentration was estimated with Qbit measurement (Invitrogen, California, USA). Prior filter-aided sample preparation (FASP) ^62^, protein extracts were incubated for 45 min with 1%v/v Benzonase Nuclease (SIGMA E1014). For the FASP digestion, the clear protein lysate was deposited in a Microcon-30kDa filter unit (Millipore, MRCF0R030) and centrifuged at 14,000g for 20 min to pass the totality of the sample through the filter. After a wash step with 200 µl buffered urea [8 M urea, 100 mM Tris-HCl (pH 8.2)] with the same centrifugation parameters as above, samples were incubated for 10 min with 50 mM iodoacetamide. The filter units were subsequently washed with 3 x 100 µl urea and 2x 100 µl 50 mM triethylammonium bicarbonate (TEAB). For overnight digestion at room temperature, sample-filters were incubated in a wet-cell adding 1:50 w/w trypsin: protein ratio (Promega, V5111) solubilized in 120 µl 50 mM TEAB. The next day, peptides were recovered by centrifugation and a small aliquot of 3 µl was taken for proteome analysis and diluted with 3% acetonitrile, 0.1% formic acid to 0.3 raw absorbance at 280 nm (NanoDrop, DeNovix).

#### Phosphopeptide enrichment

For ferric-nitrilotriacetate (Fe-NTA) phosphoenrichment, tryptic peptides were first dried in an ultra-vacuum centrifugation. Then, equal amounts for each sample (600 µg based on protein estimation) were enriched by Fe-NTA spin columns (Thermo Fisher Scientific, A32992) following the provided instructions. Dried phosphopeptides were re-dissolved in 10 µl 3% acetonitrile, 0.1% formic acid for mass spectrometry (MS) injections.

#### Mass spectrometry

MS analyses were performed on an ACQUITY UPLC M-Class System (waters) coupled to an Orbitrap Fusion Lumos (Thermo Fisher). For LC-MS/MS injections of proteome and phosphoproteome samples, 2 µl and 4 µl, respectively, were loaded on a trap column [nanoEase M/Z Symmetry C18 100 A, 5um, 1/PK 180 um x 20 mm (waters)] and separated on the analytical column [nanoEase M/Z HSS C18 T3 Col 100 A, 1.8 um, 1 P/K 75 um x 250 mm (waters)] using a linear gradient of 5-24% - 24-36% acetonitrile (0.1% formic acid) over 90 min (80 min – 10 min step) followed by an 20 min re-equilibration time. The analytical column was temperature controlled at 50°C and the flow rate of 300 nL/min was constant over the chromatographic separation.

MS survey scan from *m/z* 300-1500 were acquired at a resolution of 120,000 at *m/z* 200 in profile mode, while tandem mass spectra were acquired in the IonTrap in centroid mode. For HCD fragmentation, a normalized collision energy of 35 was applied and isolation windows of *m/z* 0.8 (proteome) or *m/z* 1.2 (phosphoproteome) were used for MS2 acquisitions. The AGC target was set to 10e5 for both types of samples, whereas the maximum injection time was either 50 ms (proteome) or 120 ms (phosphoproteome). The mass spectrometry proteomics data have been deposited to the ProteomeXchange Consortium via the PRIDE ^63^ partner repository with the dataset identifier PXD025124.

#### Protein identification and quantification

Peptide and protein identification/quantification were performed using label free quantification and match between run option in MaxQuant (version 1.6.2.3). Spectra were searched against the Uniprot *Staphylococcus aureus* reference database (downloaded 20201127), searching for the variable modifications Acetylation (Protein N-term), Oxidation (M) and if applicable Phosphorylation (STY) at 1% false discovery rate. For the proteome analyses, protein intensity estimates were used as reported in the proteinGroups.txt file. For quantification, only protein groups with at least two peptides were kept. For the phosphoproteome, the Maxquant output reported in the “Phospho STY Sites.txt” was used. Intensities were normalized to systematically remove variability in abundance due to different sample loads on the LC column. The log_2_ transformed intensities were z-scored, updated by the average of the standard deviation in all samples. After normalization, all samples had a similar intensity distribution for computing a moderated t-test ^64^, employing the R package limma ^65^. All these functions are implemented in the R package SRMService ^66^. Pathway information of proteins was retrieved from the Kyoto Encyclopedia of Genes and Genomes (KEGG) database ^67^ for visualization purposes.

#### Immunoblotting

Bacterial total protein was isolated by bacterial lysis using glass beads. Briefly, bacteria were washed twice with ice-cold PBS and resuspended in ice-cold PBS supplemented with 1X Halt Protease and Phosphatase Inhibitor (Thermo Fisher Scientific). Bacteria were mechanically lysed with glass beads in a tissue lyser (QIAGEN) for 3 cycles of 1.5 min, running at 30 Hz. Between the cycles, samples were cooled down on ice. Cell debris and intact cells were removed by centrifugation (1,200 g for 10 min at 4°C). Protein concentration was determined by Bradford assay ^68^. 10 µg of total protein were separated by sodium dodecyl sulphate polyacrylamide gel electrophoresis (SDS-PAGE) (10% acrylamide) and transferred onto a nitrocellulose membrane. Membranes were stained for total protein using the Novex™ Reversible Membrane Protein Stain Kit (Invitrogen) following the manufacturer’s instructions. After de-staining, membranes were blocked using 4% bovine serum albumin (BSA, Sigma) in Tris-buffered saline for 2 h at room temperature. An antibody recognizing phospho-Threonine (Cell Signaling, Phospho-Threonine (42H4) Mouse mAb #9386) was used to detect Thr-phosphorlyation and a polyclonal rabbit serum was used to detect the translational elongation factor EF-G, both from *S. aureus* and *B. subtilis*. Visualizing was performed using IRDye secondary antibodies (LI-COR) and the Fusion FX6 EDGE Imaging System (Vilber Lourmat S.A.).

### Statistical analysis

All experiments, except proteomics data, were analysed for statistical significance by either the Kruskal-Wallis test with Dunn’s post-test, by one-way or two-way ANOVA with Tukey post-test after normality testing by the Shapiro-Wilk normality test, unless otherwise indicated. P-values are shown were appropriate. Analysis of proteomics data is explained in the mass spectrometry section.

## Data availability

Raw data have been deposited in PRIDE ^63^: PXD025124 and are publicly available. All other study data are included in this article and supporting information.

## Acknowledgement

We thank the members of the Zinkernagel laboratory, especially Dr. Federica Andreoni and Sanne Hertegonne, for critical reading of the manuscript, Mathilde Boumasmoud for assistance with sequencing and Simon Diez (Columbia University, USA) for technical advice. Proteomic analysis was performed at the Functional Genomics Centre Zurich, University of Zurich/ETH Zurich, and we thank Dr. Sibylle Pfammatter and Dr. Jonas Grossmann for technical and bioinformatical assistance.

