## Supplementary Information for "Ser/Thr phospho-regulation by PknB and Stp mediates bacterial quiescence and antibiotic persistence in *Staphylococcus aureus*"

### Supplementary figures

#### Figure S1

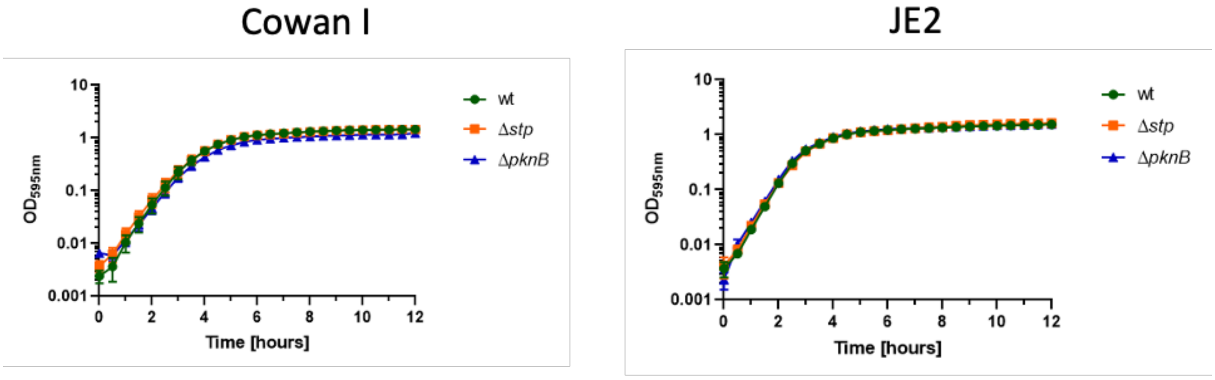


**Figure S1: Growth curves in nutrient-rich medium (TSB) show no growth defects in mutant strains.** Bacteria were grown to stationary phase in TSB shaking at 37°C for 16 hours and diluted to $OD_{600}$ 0.005 in fresh, prewarmed TSB in a 96-well flat-bottom plate (total volume 200 µl). Bacteria were incubated at 37°C and shaking 220 rpm for 12 h and $OD_{600}$ was measured every 30 min on a Tecan m200 nano plate reader. N= 3 biological replicates.

#### Figure S2

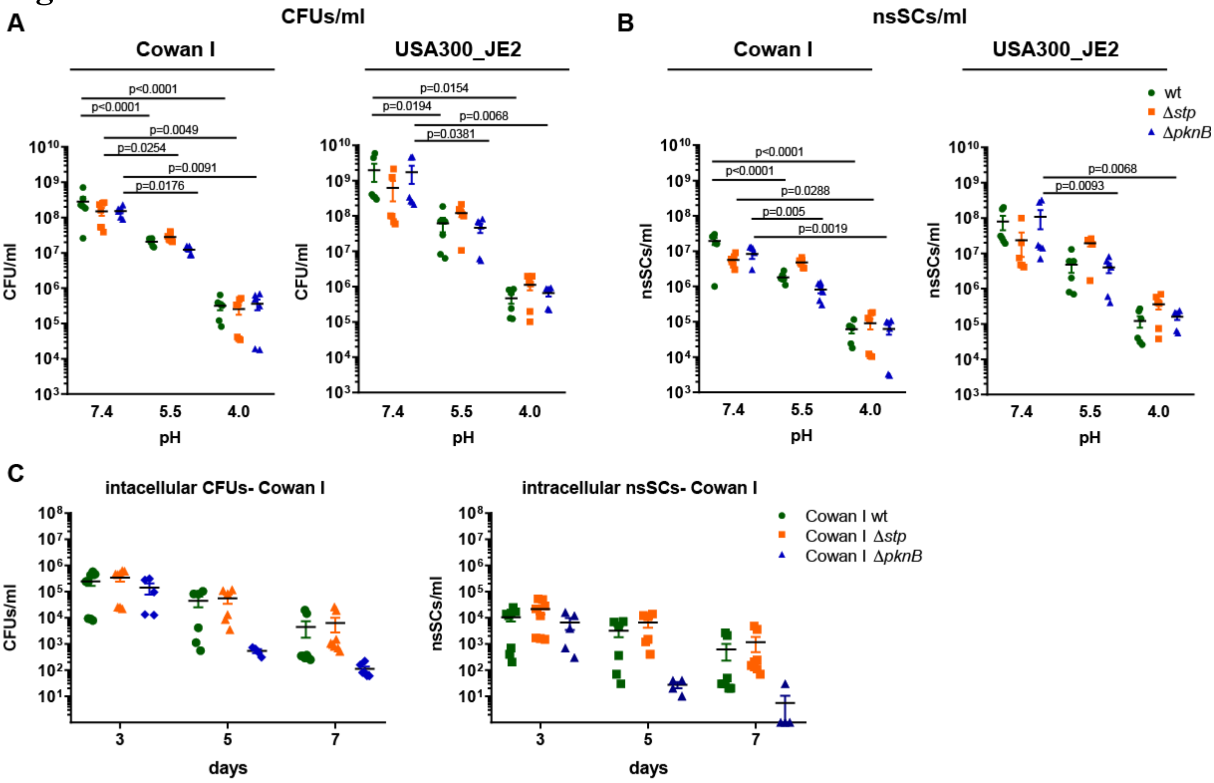


**Figure S2: Total numbers of colony forming units (CFUs) and non-stable small colonies (nsSCs) after different *in vitro* challenges. A)** Numbers of CFUs/ml after 3 days exposure to pH 7.4, pH 5.5 or pH 4.0 of *S. aureus* Cowan I and USA300_JE2 wildtype and their isogenic mutants Δ*stp* and Δ*pknB*. **B)** Numbers of nsSCs/ml after 3 days exposure to pH 7.4, pH 5.5 or pH 4.0 of *S. aureus* Cowan I and USA300_JE2 wildtype and their isogenic mutants Δ*stp* and Δ*pknB*. **C)** Numbers of CFUs and nsSCs of *S. aureus* Cowan I after exposure to the intracellular milieu of A549 cells for 3, 5 or 7 days. Statistical significance was determined using two-way ANOVA with Tukey post-test.

#### Figure S3

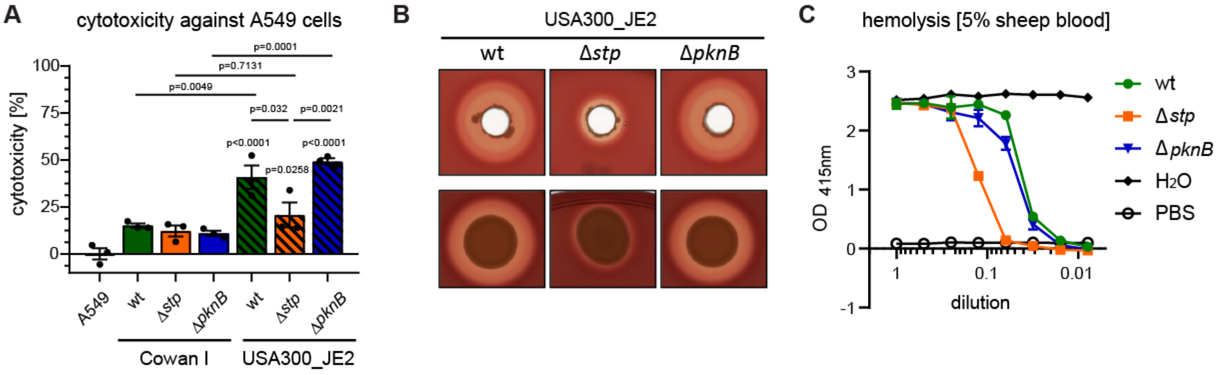


**Figure S3: Cytotoxicity and haemolytic activity of *S. aureus* wt, Δ*stp* and Δ*pknB* mutants. A)** Assessment of bacterial cytotoxicity towards the lung epithelial cell line A549 by MTT assay. No differences in cytotoxicity were observed for Cowan I and its isogenic mutants Δ*stp* and Δ*pknB* which in general showed low levels of cytotoxicity. USA300_JE2 wildtype and Δ*pknB* showed the highest levels of cytotoxicity with almost 50% cell death after 24 h infection (eradication of extracellular bacteria 3.5 h post infection). USA300_JE2 Δ*stp* displayed a significantly reduced cytotoxicity as compared with the wildtype or Δ*pknB*. Statistical significance was determined using one-way ANOVA with Tukey post-test. **B)** Haemolytic activity is reduced in USA300_JE2 Δ*stp*. Bacteria were grown to stationary phase in Todd Hewitt (TH) medium at 37°C and shaking 220 rpm for 16 h. 10 µl of stationary phase culture were inoculated in 5 mm punch holes (*upper panel*) or 25 µl directly on a Columbia agar plate supplemented with 5% sheep blood (BioMérieux) (*lower panel*) and incubated for 24 h at 37°C and 24 h at 4°C. **C)** Quantification of haemolytic activity confirmed the reduced haemolytic activity of USA300_JE2 Δ*stp*.

#### Figure S4

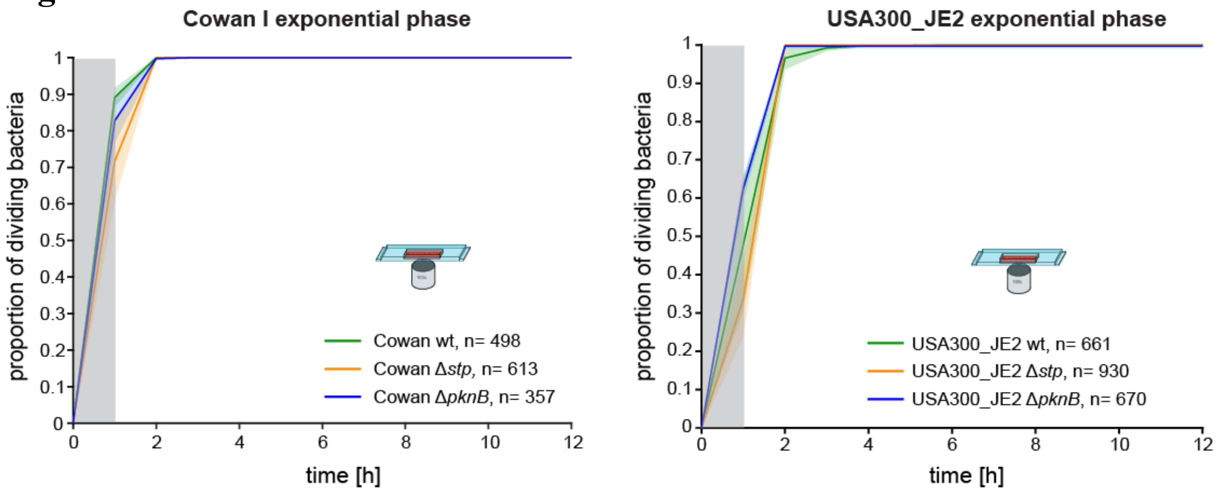


**Figure S4: Single cell microscopy of exponentially growing bacteria shows no growth difference between wildtype and isogenic mutants.** Bacteria were grown to exponential phase in TSB at 37°C and shaking 220 rpm for 2 h before imaging by time-lapse microscopy. Grey zone marks the period at the beginning of the experiment, where cell divisions could occur, but would not be observed.

### Figure S5

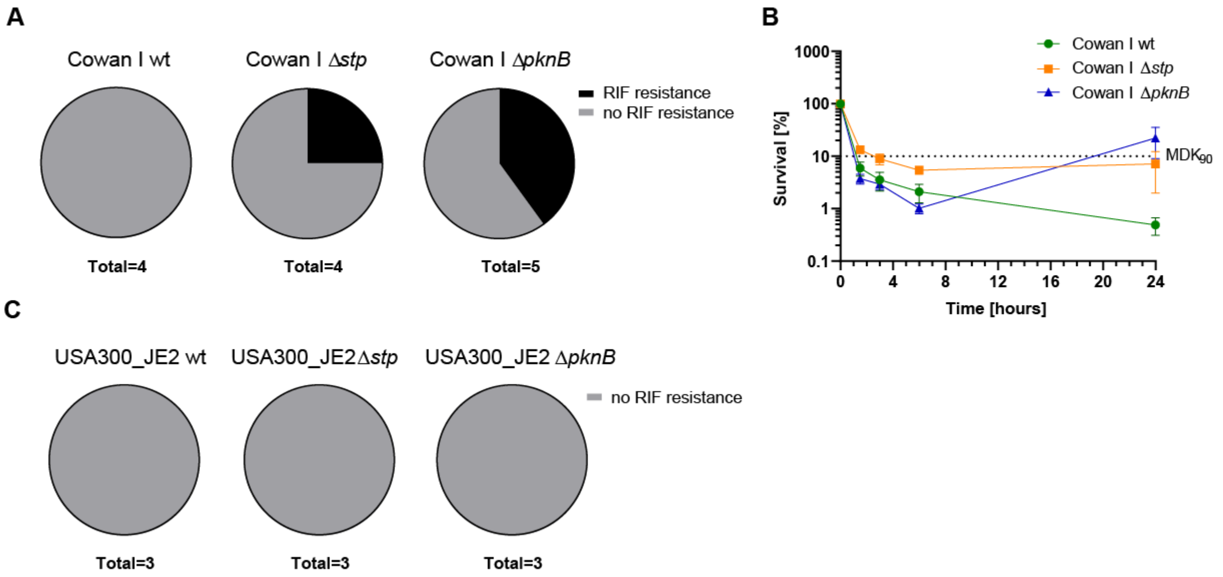


**Figure S5: Rifampicin resistance development during time kill assays in *S. aureus*. A)** Pie charts show number of events where resistance to rifampicin developed during the experiment in Cowan I. Experiments where resistance developed where excluded from Fig. 3. **B)** Time kill curves for Cowan I wildtype and its isogenic mutants Δ*stp* and Δ*pknB* in 40x minimum inhibitory concentration (40xMIC) of rifampicin including experiments with RIF-resistance development. **C)** Pie charts show that no resistance to rifampicin developed during the experiment in USA300_JE2.

#### Figure S6

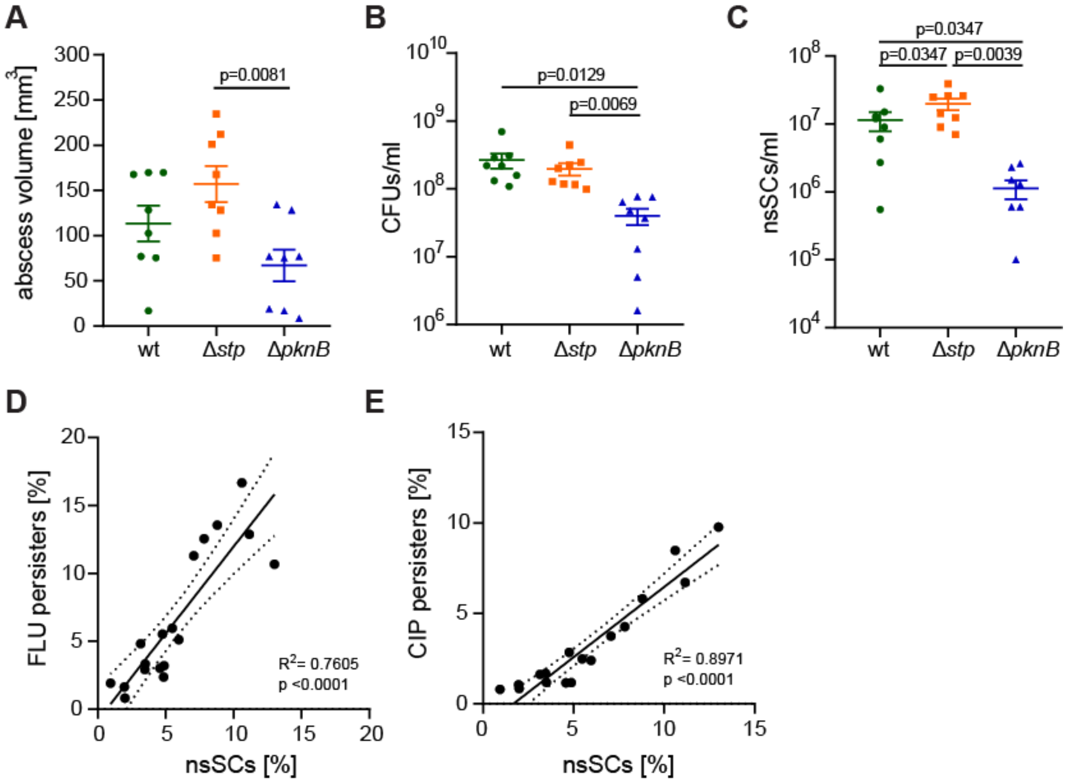


**Figure S6: Murine infection model shows increased abscess volume and non-stable small colony formation in *S. aureus* Cowan I Δ*stp*. A)** Calculated abscess volume at day 5 post infection with Cowan I wildtype, Δ*stp* and Δ*pknB*. **B)** Bacterial load in murine abscess pus five days post infection. **C)** Number of nsSCs in abscess pus five days p.i. **A-C)** Statistical significance was determined by one-way ANOVA with Tukey post-test. **D)** Linear regression analysis on FLU persisters (%) and nsSCs (%) in murine abscess pus five days p.i. Dots represent values from individual abscesses from mice infected with *S. aureus* Cowan I wildtype, Δ*pknB* or Δ*stp* five days p.i. **E)** Linear regression analysis on CIP persisters (%) and nsSCs (%) in murine abscess pus five days p.i. Dots represent values from individual abscesses from mice infected with *S. aureus* Cowan I wildtype, Δ*pknB* or Δ*stp* five days p.i. N= 6 abscesses from 3 mice.

#### Figure S7

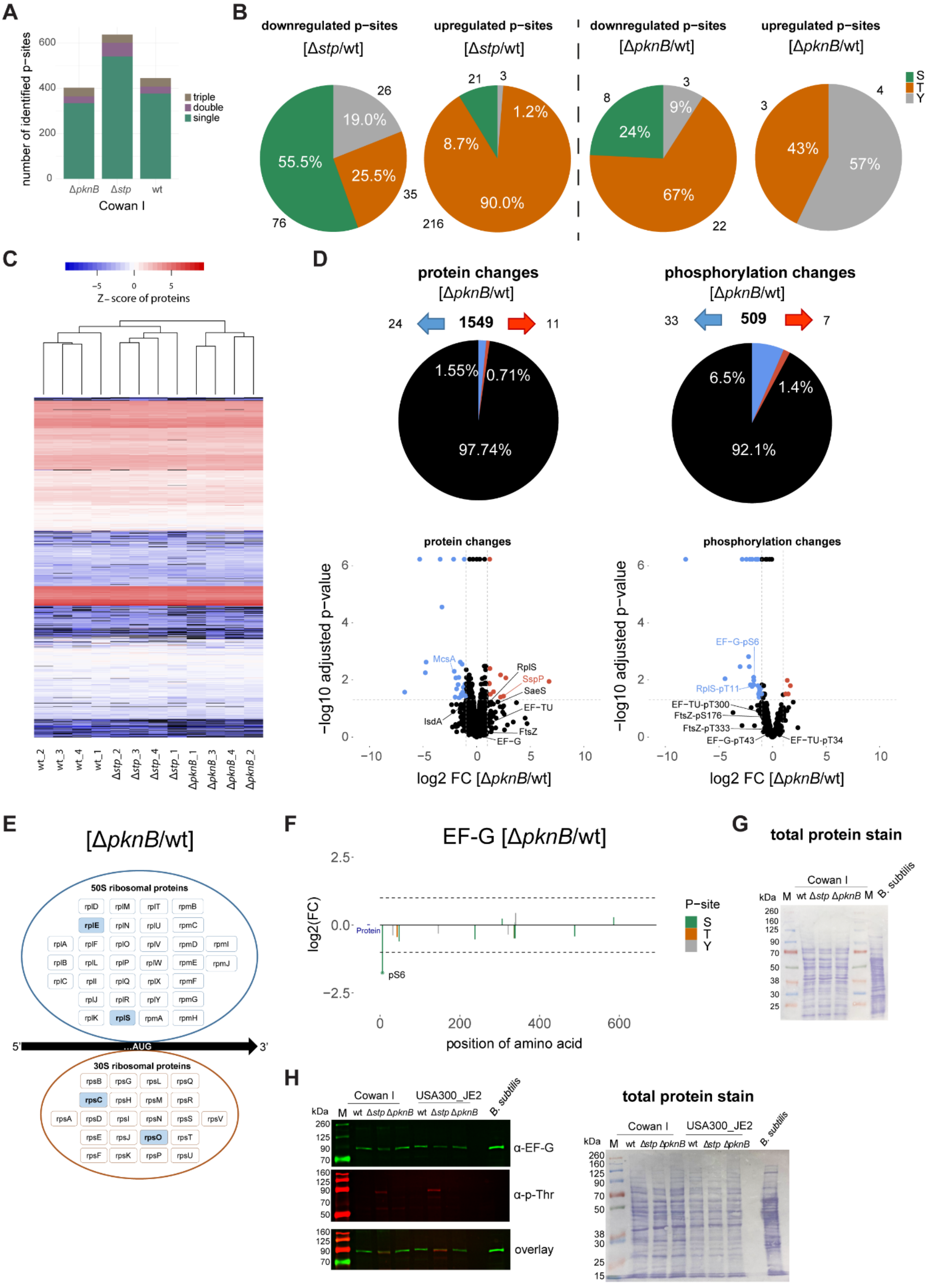


**Figure S7: Proteomic and phosphoproteomic analysis of pH 5.5 exposed *S. aureus* Cowan I. A)** Number of identified phosphorylation sites and their multiplicity in Cowan I wildtype, Δ*stp* and Δ*pknB* after pH 5.5

exposure for three days. **B)** Pie charts showing up- and downregulated phosphorylation sites (Ser, Thr, Tyr) for Δ*stp* and Δ*pknB* in comparison to wildtype. **C)** Hierarchical cluster analysis of protein abundance in proteome. The abundance profiles of the proteins in our four replicates hierarchically clustered by heatmap. Blue indicates less abundant proteins, whereas red indicates more abundant proteins. Black indicates non-detected proteins. **D)** Pie chart and volcano plot comparing proteins between Cowan I *pknB* deletion mutant and wildtype, identifying 35 differentially expressed proteins. Twenty-four proteins were underrepresented (blue symbols) and 11 proteins were overrepresented (red symbols) (*left panel*). Pie chart and volcano plot comparing phosphopeptides between Cowan I *pknB* deletion mutant and wildtype, identifying 40 differentially expressed phosphopeptides. Thirty-three phosphopeptides were underrepresented (blue symbols) and seven phosphopeptides were overrepresented (red symbols) (*right panel*). Cut-off adjusted p-value < 0.05 and a fold change of |log2FC| > 1. Underrepresented proteins/ phosphopeptides are depicted by blue symbols and overrepresented proteins/ phosphopeptides with red symbols. **E)** Blue colour indicates decreased Ser/Thr phosphorylation (at least one phospho-site log2 ≦ -1, adjusted p-value < 0.05). If upregulated and downregulated phosphopeptides for the same protein were found, the highest up- or downregulation is shown. **F)** p-sites to protein plot for the elongation factor EFG. Phosphorylation sites and exact amino acid positions are indicated. Y-axis indicates log2 fold changes of phosphopeptides in Cowan I Δ*pknB* compared to wildtype. **G)** Total protein stain as loading control for immunoblot in Fig. 5F. **H)** Representative immunoblot of bacterial cell lysates with antibody specific for phospho-Thr and rabbit serum raised against *B. subtilis* EF-G that also recognises *S. aureus* EF-G. Cowan I and USA300_JE2 wildtype, Δ*stp* and Δ*pknB* were grown at pH 5.5 for three days, *B. subtilis* was grown to stationary phase for 16 h.

### Figure S8

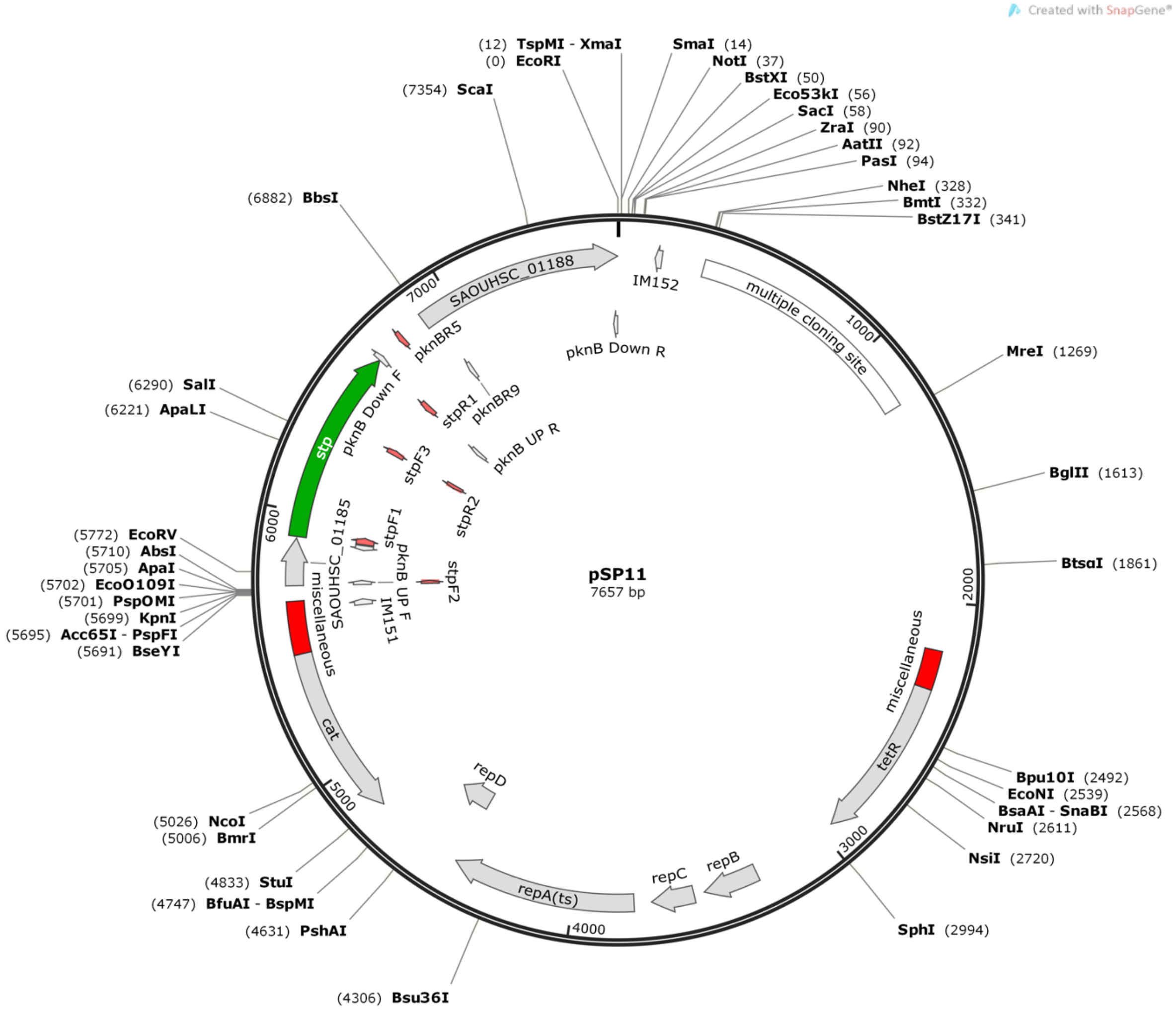


**Figure S8: Plasmid map of pSP11.** The pIMAY plasmid was used as backbone modified accordingly to delete *pknB* by homologous recombination in *S. aureus*.

### Figure S9

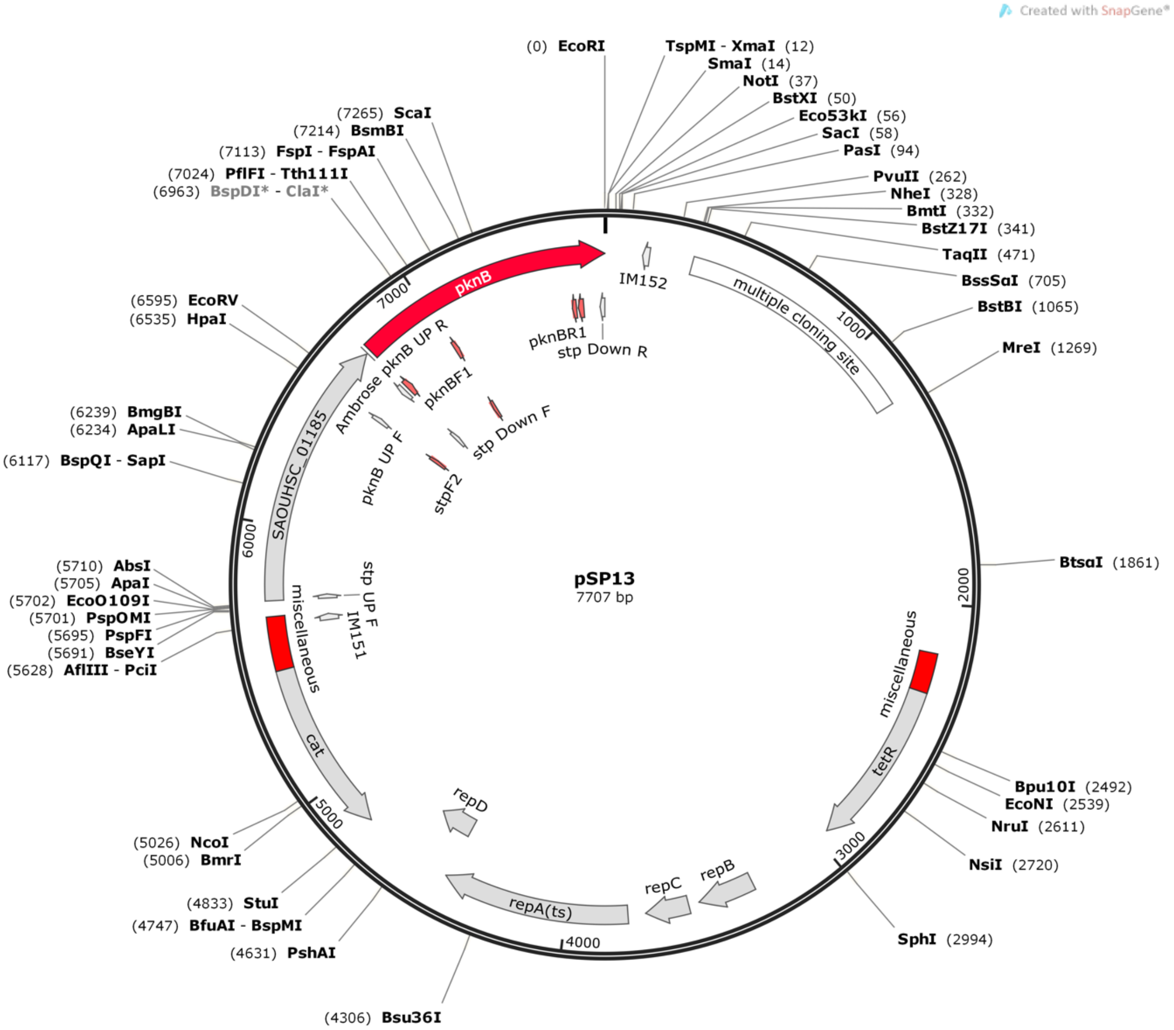


**Figure S9: Plasmid map of pSP13.** The pIMAY plasmid was used as backbone modified accordingly to delete *stp* by homologous recombination in *S. aureus*.

#### Figure S10

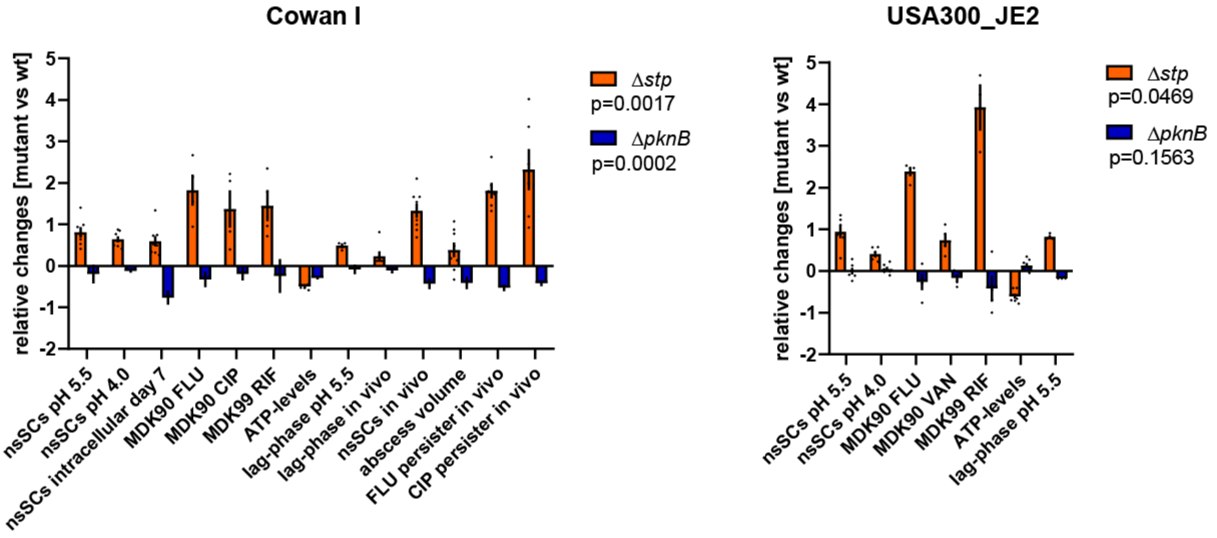


**Figure S10: Summary of the individual assay results in relation to the wildtype for *S. aureus* Cowan I and USA300_JE2.** Dots represent individual biological replicates. Deletion of *stp* leads to increased proportions of non-stable small colonies (nsSCs), prolonged lag-phases, elevated antibiotic persistence and reduced ATP levels in *S. aureus* Cowan I and USA300_JE2 after acid stress exposure. An *in vivo* murine abscess environment results in prolonged lag-phases, an increased proportion of nsSCs, larger abscesses and high levels of antibiotic persistence in the *stp* deletion mutant of Cowan I. An opposite behaviour was observed for the *pknB* deletion mutants. Statistically significant deviation from wildtype was calculated by Wilcoxon matched-pairs signed rank test.

#### Supplementary tables

##### Table S1: Bacterial strains

| Strain | Description | Reference, Source |
|---|---|---|
| *E. coli* DC10B | universal Staphylococcal clonal host | Invitrogen |
| *S. aureus* Cowan I wt | MSSA, ST30, wildtype | ATCC 12598 |
| *S. aureus* Cowan I Δ*stp* | MSSA, ST30, *stp* deletion | this study |
| *S. aureus* Cowan I Δ*pknB* | MSSA, ST30, *pknB* deletion | this study |
| *S. aureus* USA300_JE2 wt | MRSA, ST8, wildtype | NARSA Collection |
| *S. aureus* USA300_JE2 Δ*stp* | MRSA, ST8, *stp* deletion | this study |
| *S. aureus* USA300_JE2 Δ*pknB* | MRSA, ST8, *pknB* deletion | this study |

##### Table S2: Primers and plasmids used for deletion of *pknB* and *stp*.

| Primers | Sequence 5´→ 3´ | Function | Reference |
|---|---|---|---|
| pknB UP F2 | gatcccccgggctgcaggaattCCTGTCAACCATGTTCCAG | Amplify Upstream pknB, fuse to Downstream pknB and clone in pIMAY by Gibson assembly | this study |
| pknB UP R2 | tttacttcaaTCATACTTTATCACCTTCAATAGC | Amplify Upstream pknB, fuse to Downstream pknB and clone in pIMAY by Gibson assembly | this study |
| pknB DOWN F2 | gaacaaaagctgggtaccGCTTAACATTACAATTAGGTTCTTTG | Amplify Downstream pknB, fuse to Upstream pknB and clone in pIMAY by Gibson assembly | this study |
| pknB DOWN R2 | gaacaaaagctgggtaccGCTTAACATTACAATTAGGTTCTTTG | Amplify Downstream pknB, fuse to Upstream pknB and clone in pIMAY by Gibson assembly | this study |
| stp UP F | tagtctcgagCTGGCTCGTTGAACAAGG | Amplify UPstream region of stp for KO | this study |
| stp UP R | tatcaccttcGTCTTTACCTCGTTTCTACTTG | Amplify UPstream region of stp for KO | this study |
| stp DOWN F | aggtaaagacGAAGGTGATAAAGTATGATAGG | Amplify Downstream region of stp for KO | this study |
| stp DOWN R | ctctgaattCCATTTACAATAGGTACTTGC | Amplify Downstream region of stp for KO | this study |
| stp UP R2 | tttacttcaaGTCTTTACCTCGTTTCTACTTG | Amplify UPstream region of stp for KO of both stp and pknB | this study |
| IM151 | TACATGTCAAGAATAAACTGCCAAAGC | Check for presence of pIMAY | [1] |
| IM152 | AATACCTGTGACGGAAGATCACTTCG | Check for presence of pIMAY | [1] |
| **Plasmids** | **-** | **Function** | **Reference** |
| pIMAY | - | Backbone plasmid | [1] |
| pSP11 | - | Plasmid for *pknB* KO | this study |
| pSP13 | - | Plasmid for *stp* KO | this study |

**Table S3: Minimum Inhibitory Concentrations (MICs) for different antibiotics in DMEM +10% FCS +1% L-Glu pH 7.4.**

| MICs (mg/L) | Cowan I | | | USA300_JE2 | | |
|---|---|---|---|---|---|---|
| **Antibiotic** | **wt** | **Δstp** | **ΔpknB** | **wt** | **Δstp** | **ΔpknB** |
| Flucloxacillin (FLU) | 0.25 | 0.5 | 0.25 | 0.5 | 1 | 0.5 |
| Ciprofloxacin (CIP) | 0.008 | 0.008 | 0.008 | 16 | 16 | 16 |
| Vancomycin (VAN) | nd | nd | nd | 1 | 2 | 1 |
| Rifampicin (RIF) | 1 | 1 | 1 | 0.015 | 0.015 | 0.015 |

### Material and Methods

#### Quantitative haemolysis assay.

Overnight cultures of USA300_JE2 and its corresponding mutants were grown in Todd Hewitt (TH) medium as previously described [2,3]. Optical density (OD) was adjusted to $OD_{600}$ 2 and bacteria were centrifuged, supernatant sterile filtered and 100 µl were added to 100 µl of washed 5% sheep blood (Thermo Fisher). After incubation at 37°C for 30 min and 4°C for 30 min, the suspension was centrifuged at 1,600 rpm for 15 min and haemoglobin absorbance was measured in the supernatant at 415 nm with the VERSAmax tuneable microplate reader (Molecular Device).

#### Cell viability assay

To assess bacterial cytotoxicity, a MTT assay was performed. As previously described [4], epithelial cells (A549, ATCC CCL-185) were infected at a multiplicity of infection (MOI) of 1. After 3.5 h, extracellular bacteria were inactivated by addition of flucloxacillin (Actavis, 1 mg/ml) and lysostaphin (Sigma, 25 µg/ml) (invasion). Twenty-four hours after invasion, the tetrazolium dye MTT (3-[4,5-dimethylthiazole-2-yl]-2,5-diphenyltetrazolium bromide, Sigma) was added to the medium to a final concentration of 1 mg/ml and incubated for 2 h at 37°C and 5% $CO_2$. After removing the medium, the formed formazan crystals were dissolved in 0.04 M HCl in isopropanol. Absorbance was measured at 570 nm with the VERSAmax tuneable microplate reader (Molecular Device). Percentage of survival of the eukaryotic cells was calculated relative to an uninfected control.
